# Structural basis for growth factor and nutrient signal integration on the lysosomal membrane by mTORC1

**DOI:** 10.1101/2024.11.15.623810

**Authors:** Zhicheng Cui, Alessandra Esposito, Gennaro Napolitano, Andrea Ballabio, James H. Hurley

## Abstract

Mechanistic target of rapamycin complex 1 (mTORC1), which consists of mTOR, Raptor, and mLST8, receives signaling inputs from growth factor signals and nutrients. These signals are mediated by the Rheb and Rag small GTPases, respectively, which activate mTORC1 on the cytosolic face of the lysosome membrane. We biochemically reconstituted the activation of mTORC1 on membranes by physiological submicromolar concentrations of Rheb, Rags, and Ragulator. We determined the cryo-EM structure and found that Raptor and mTOR directly interact with the membrane at anchor points separated by up to 230 Å across the membrane surface. Full engagement of the membrane anchors is required for maximal activation, which is brought about by alignment of the catalytic residues in the mTOR kinase active site. The observations show at the molecular and atomic scale how converging signals from growth factors and nutrients drive mTORC1 recruitment to and activation on the lysosomal membrane in a three-step process, consisting of (1) Rag-Ragulator-driven recruitment to within ∼100 Å of the lysosomal membrane, (2) Rheb-driven recruitment to within ∼40 Å, and finally (3) direct engagement of mTOR and Raptor with the membrane. The combination of Rheb and membrane engagement leads to full catalytic activation, providing a structural explanation for growth factor and nutrient signal integration at the lysosome.

## Main

The mechanistic target of rapamycin complex 1 (mTORC1) integrates upstream growth factor and nutrient signals to stimulate anabolic processes connected to cell growth and inhibit catabolic processes such as autophagy ^1,2^. Growth factors (GFs) acting on G-protein coupled receptors (GPCRs) and receptor tyrosine kinases (RTKs) at the plasma membrane (PM) initiate signaling through class I phosphoinositide 3-kinase (PI3K), the PIP_3_-activated protein kinase Akt, the tuberous sclerosis complex (TSC) complex. TSC in turn regulates the lysosomally-localized small GTPase Ras homologue enriched in brain (Rheb) ^3^. Direct binding of Rheb-GTP to the mTOR kinase subunit of mTORC1 allosterically activates the kinase by inducing a large-scale conformational change ^4^. Rheb is required for phosphorylation of the canonical substrates of mTORC1 ^5^, including the ribosomal protein S6 kinase (S6K) and eukaryotic translation initiation factor 4E binding protein-1 (4EBP1) that mediate mTORC1 stimulation of protein synthesis ^1^, whereas it is dispensable for the phosphorylation of non-canonical substrates, such as transcription factor EB (TFEB) ^6^. Rheb is present in cells at an estimated 650 nM ^7^, yet > 100 μM soluble Rheb-GTP is needed for half-maximal mTORC1 activation ^4^. Thus, there is a greater than two orders of magnitude discrepancy between the physiological levels and the *in vitro* biochemistry of Rheb, the most fundamental activator of mTORC1.

Rheb is farnesylated ^8–10^, which is essential for mTORC1 activation and is responsible for targeting Rheb to lysosomes. Despite that the receptor-PI3K-Akt signaling pathway originates at the PM, Rheb signaling to mTORC1 occurs exclusively on the cytosolic face of lysosomes ^3^. mTORC1 is recruited to lysosomes by the Ras-related GTP binding (Rag) GTPases RagA-D under amino acid replete conditions ^11–13^. The Rags function as heterodimers, with RagA or RagB paired with RagC or RagD. Active Rag dimers consisting of RagA/B-GTP and RagC/D-GDP recruit mTORC1 to the lysosome by binding to its Raptor subunit ^14,15^. The Rags, in turn, are recruited to the lysosome membrane by the pentameric Ragulator complex, specifically, by its myristoylated and palmitoylated Lamtor1 subunit ^16^. Amino acid-dependent mTORC1 recruitment to lysosomes by the Rags and Ragulator brings mTORC1 into proximity to the lysosome-bound pool of Rheb-GTP. This serves as a physiological “AND” gate for a GF signal and an ample pool of biosynthetic precursors prior to protein synthesis and cell growth. How this physiologically critical AND gate might be organized and implemented at the structural level is unknown.

We hypothesized that the lysosome membrane itself might be the missing link that orchestrates the AND gate and contributes the missing > 2 orders of magnitude needed for Rheb-GTP activation of mTORC1. The feasibility of atomistic single-particle cryo-EM reconstructions of liposome-bound peripheral protein assemblies has recently been demonstrated ^17,18^. This prompted us to reconstitute the concerted activation of mTORC1 on liposomes by Rheb and RagA/C-Ragulator, and elucidate its structural basis.

### Reconstitution of mTORC1 activation on membranes

We employed Large Unilamellar Vesicles (LUVs) with a lipid composition of 72.8% DOPC, 7% POPS, 10% Cholesterol, 5% DGS-NTA, 5% PE-MCC, and 0.2% DiR, as the membrane platform to investigate the role of Rag-Ragulator and Rheb in mTORC1 activation. The active RagA-GTP-RagC-GDP-Ragulator complex was recruited to LUVs containing the lipid DGS-NTA(Ni) via a 6xHis-tag fused to the N-terminus of the Lamtor1 subunit of Ragulator in place of the physiological myristoyl and palmitoyl modifications (Fig. 1a). Rheb was tethered to the LUVs via a thiol-maleimide reaction between the functionalized lipid PE-MCC and its only cysteine residue, C181, which is the farnesylation site of endogenous Rheb (Fig. 1a). Rheb and RagA/C-Ragulator were used at essentially physiological concentrations of 250 and 300 nM, respectively. We monitored mTORC1 kinase activity as a function of LUVs, Rheb, and Rag-Ragulator by detecting the phosphorylation of Thr37 and Thr46 of full-length 4EBP1. mTORC1 kinase activity increased over 35-fold in the presence of LUVs, Rag-Ragulator, and Rheb-GTP (Fig. 1b), but not with Rheb-GDP. As expected, no activity was observed in the presence of the mTOR inhibitor Torin1 or the absence of mTORC1. LUVs and Rheb-GTP increased mTORC1 activity by ∼ 3-fold, whereas other combinations did not affect its activity. No increase in mTORC1 activity was observed when 250 nM Rheb-GTP was present but LUVs were not, consistent with the past report that > 100 μM soluble Rheb-GTP is required for activation ^4^. Therefore, the combination of liposomes and liposome-tethered Rheb and RagA/C-Ragulator synergistically and potently activates mTORC1 phosphorylation of 4EBP1.

**Fig. 1:**
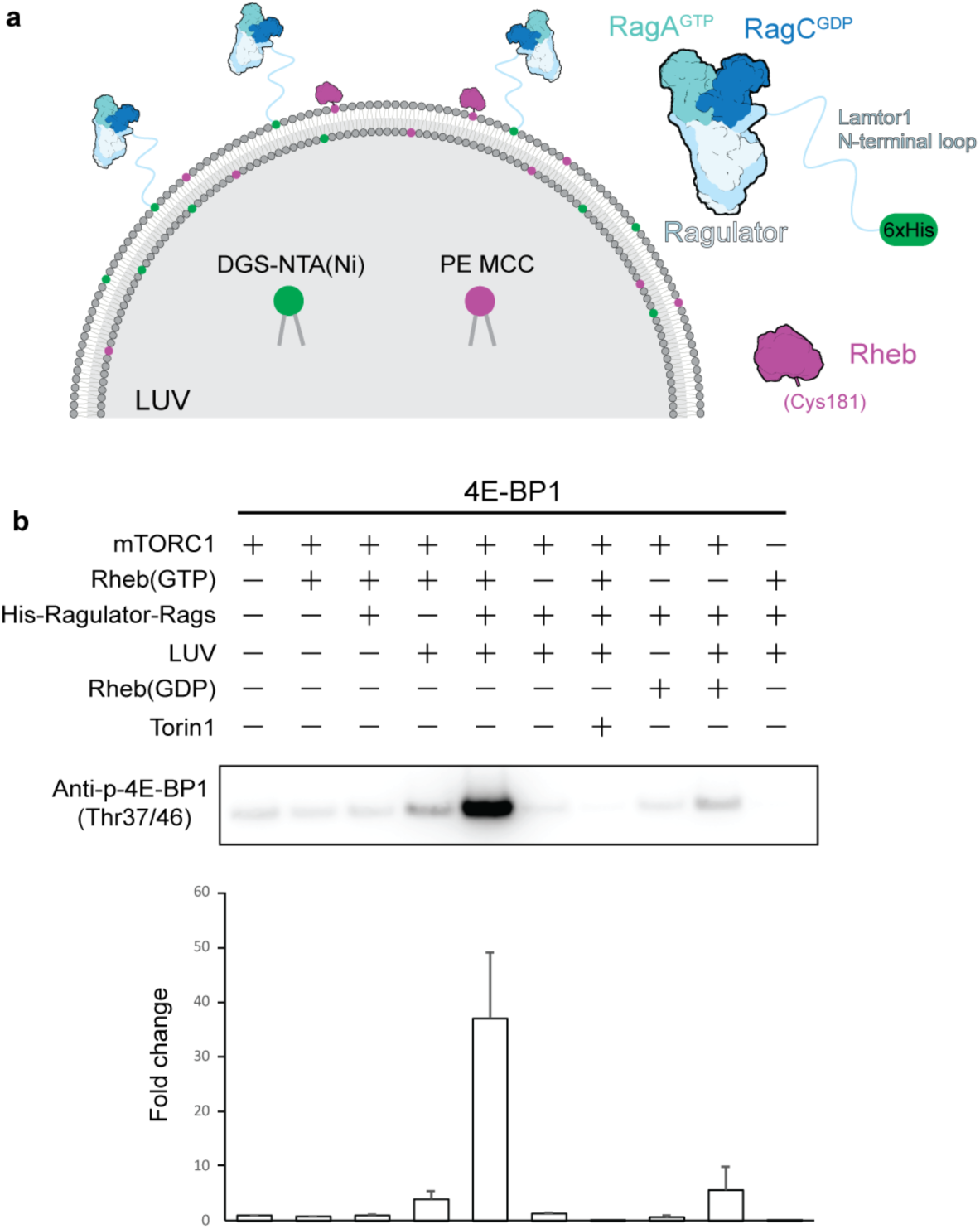
In vitro reconstitution and mTORC1 kinase activity on membrane. **a**, cartoon showing the in vitro membrane system for testing mTORC1 kinase activity. N-terminal 6xHis tag of Lamtor1 interacts with DGS-NTA(Ni) and Cys 181 of Rheb interacts with PE-MCC. **b**, Screening of components in mTORC1 activation. Western blot is done by anti-phospho-4EBP1 (Thr37/46). The quantification is shown with 3 repeats.

### Cryo-EM structure of mTORC1-Rheb-Rag-Ragulator-4EBP1 assembled on a membrane

We reconstituted the mTORC1-Rheb-Rag-Ragulator with the full-length substrate 4EBP1 complex on liposomes, and acquired cryo-EM images (Fig. 2a). 2D class averages suggested that the complex was rigidly oriented with respect to the phospholipid bilayer (Fig. 2b). We determined the cryo-EM structure of the entire assembly to an overall resolution of 3.23 Å (Extended Data Fig. 1). The overall structure of mTORC1 and its interactions with Rheb and Rag-Ragulator are consistent with previous structures determined in the absence of membranes ^4,14,15,19,20^, including the large conformational change induced by Rheb binding ^4^. However, when superimposing the membrane-bound mTORC1-Rheb structure with the soluble mTORC1-Rheb structure based on the mTOR subunit, we observed a conformational change in the Raptor subunit, showing a rotation of about 7 degrees toward the HEAT domain of mTOR (Extended Data Fig. 2). This rotational movement of Raptor in the membrane-bound mTORC1-Rheb aligns with the transition from apo mTORC1 to soluble mTORC1-Rheb, suggesting that the membrane plays a role in further constricting mTORC1 when bound to Rheb. Local refinement of the mTOR-Rheb-mLST8 and Raptor-Rag-Ragulator subcomplexes yielded cryo-EM density maps at resolutions of 3.12 Å and 2.98 Å, respectively, allowing us to build atomically detailed models of the entire assembly (Fig. 3c,d). The high-resolution cryo-EM density of nucleotides confirmed the active states of Rag GTPases (Extended Data Fig. 3a,b). The density of inositol hexakisphosphate (IP6) was observed, surrounded by the lysine/Arginine cluster in the FAT domain of mTOR as previously reported in the structures of mTORC2 ^21^ and the TFEB-containing megacomplex of mTORC1 ^20^ (Fig. 3e). Despite the inclusion of full-length 4EBP1, only the TOR signaling (TOS) motif was visualized (Fig. 3f), bound to the same location on the Raptor subunit as previously identified ^4,22^.

**Fig. 2:**
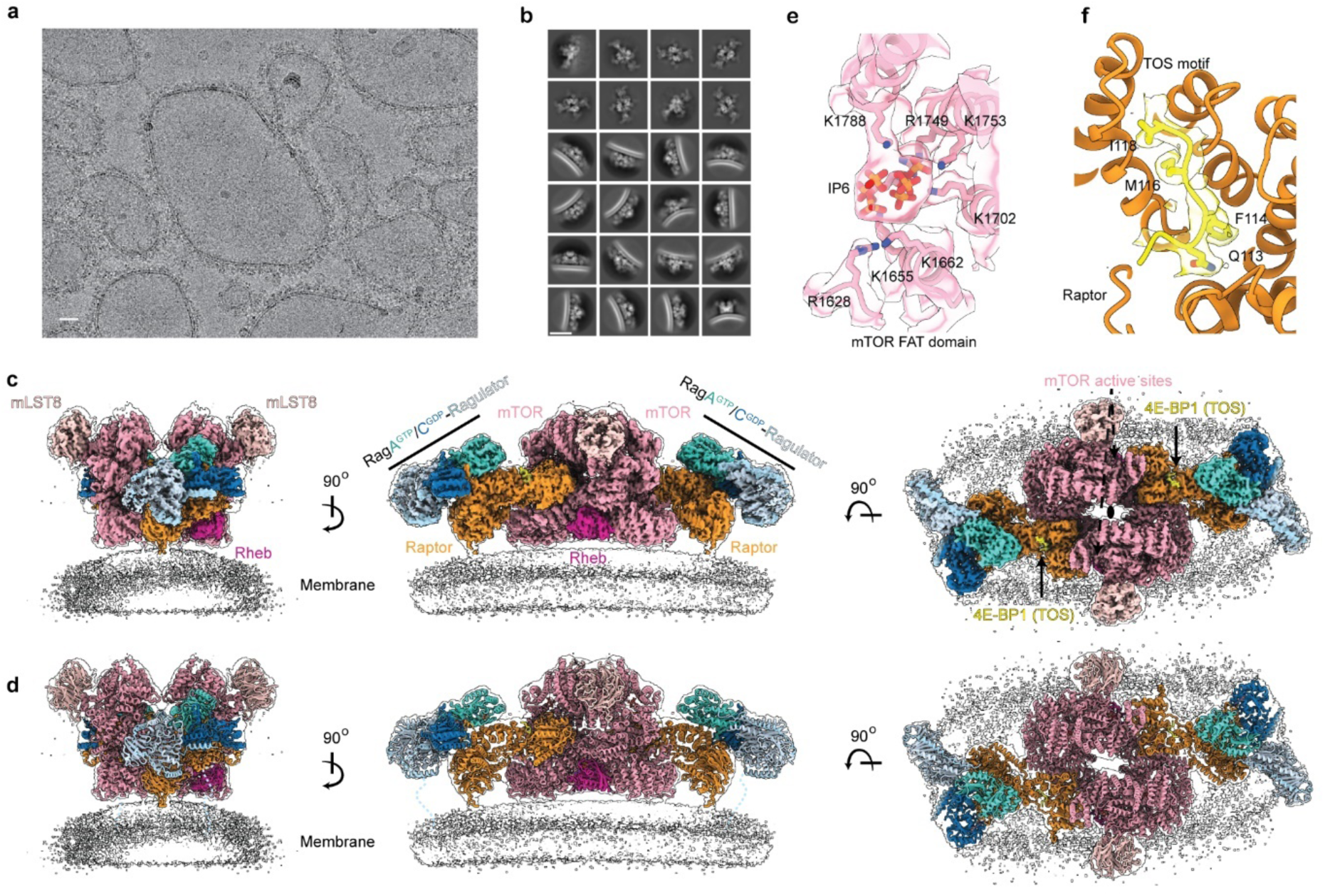
cryo-EM structure of mTORC1-Rheb-Rag-Ragulator-4EBP1 on membrane. **a**, Representative cryo-EM images showing protein-decorated liposomes. **b**, Representative 2D averages showing side and top views of the protein-membrane complex. **c**, A composite cryo-EM density map of mTORC1-Rheb-Rag-Ragulator-4EBP1 on membrane, assembled from two focused-refinement maps (mTOR-Rheb-mLST8 and Raptor-Rag–Ragulator), overlaid with the unsharpened cryo-EM map from the overall refinement with C2 symmetry. The active sites of mTOR are labeled with dashed arrows. The twofold axis is labeled as an oval symbol in the top view. Different contour levels were used for optimal visualization using UCSF ChimeraX. **d**, Atomic model of mTORC1-Rheb-Rag-Ragulator-4EBP1 is overlaid with the unsharpened cryo-EM map from the overall refinement with C2 symmetry. **e**, Close-up view of the density for inositol hexakisphosphate (IP6) and surrounding Lysin/Arginine cluster. **f**, Close-up view of the density for the 4EBP1 TOS motif. Scale bars in (**a**) and (**b**) represent 20 nm.

**Fig. 3:**
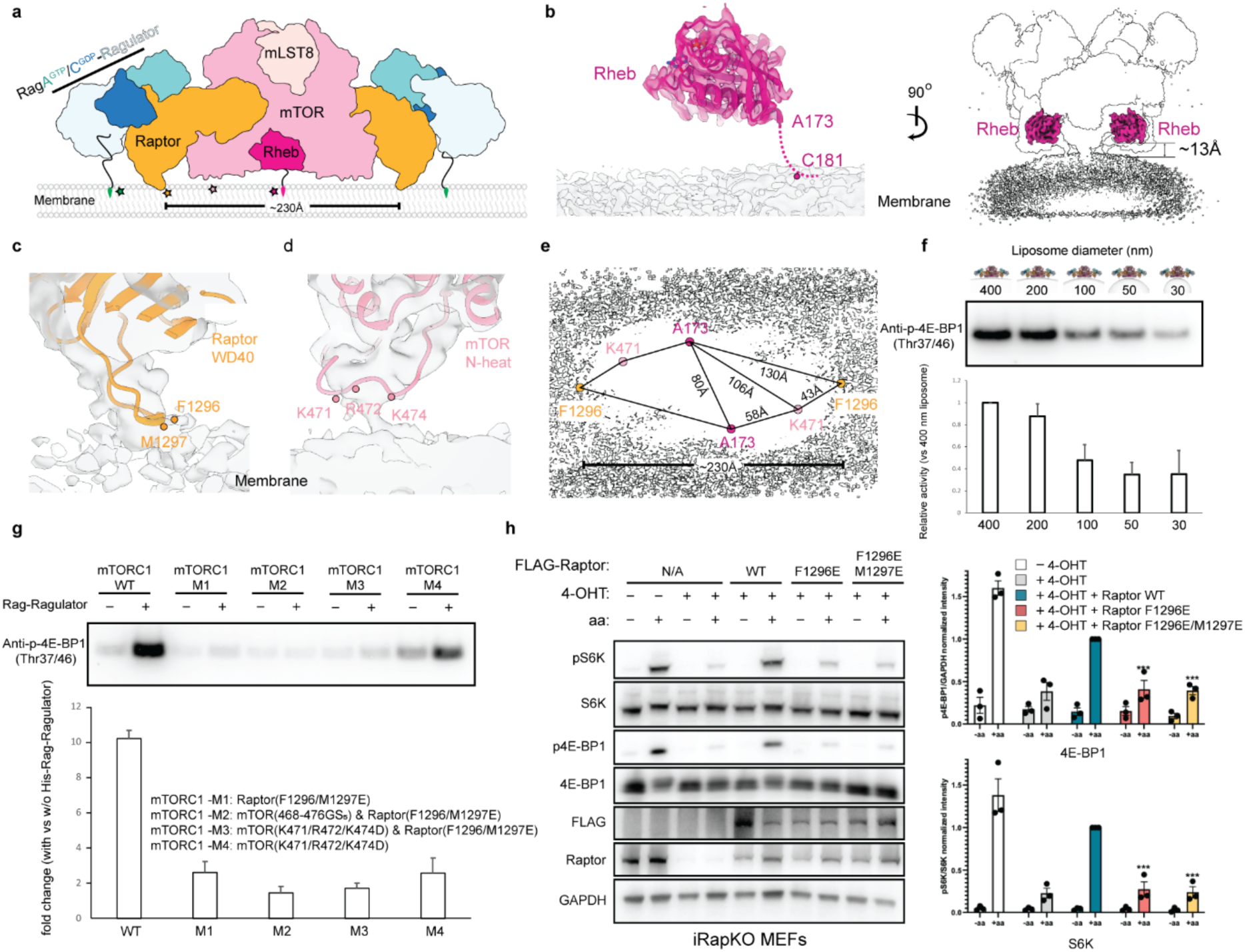
Membrane-interacting sites of mTOR and Raptor subunits. **a**, A carton representation of the mTORC1-Rheb-Rag-Ragulator-4EBP1 complex on membrane shown from the side view. The distance between two Raptor membrane-interacting sites is shown. Colored stars indicate the membrane contact sites. The lipidation sites of Lamtor1 and Rheb are indicated at the end of the arbitrary linkers that anchor them to membranes. **b**, Close-up view of Rheb density above the membrane (left), and overall Rheb position relative to the membrane (right). The last visible residue is shown, and the potential membrane-tethering loop is indicated with a dashed line. Close-up view of membrane-interacting sites of Raptor (**c**) and mTOR (**d**) subunits. For better visualization, the unsharpened cryo-EM map from the overall refinement with C2 symmetry is used. Residues of Raptor and mTOR are indicated in dots. **e**, The geometry of the residues involved in membrane interaction. **f**, In vitro kinase activity of mTORC1 kinase activity with different liposome sizes, the quantification is done with 3 repeats. **g**, In vitro kinase activity of mTORC1 mutants, the quantification is shown with 3 or 4 repeats. **h**, The Inducible Raptor KO (iRapKO) MEFs, treated with 0.5 μM 4-hydroxy-tamoxifen (4-OHT) for 48h or left untreated, were transfected with either WT, F1296E, or F1296E/M1297E Raptor mutants. 24h after transfection, cells were starved of amino acids for 60 min or starved and re-stimulated with amino acids for 30 min and analysed by immunoblotting with the indicated antibodies. Quantifications of pS6K/S6K and p4EBP1/GAPDH are shown with mean ± s.e. throughout; n = 3 experiments (***P < 0.0001, two-way ANOVA, Dunnett’s multiple comparisons test).

We noticed two 3D classes with extra Rag-Ragulator densities bound to the mLST8 subunit of mTORC1, and refined to overall resolutions of 3.47 Å and 3.81 Å for classes with the one and two extra copies of Rag-Ragulator, respectively (Extended Data Fig. 4). We pooled these classes together and carried out a focused refinement of the mLST8-Rag-Ragulator subcomplex, yielding a cryo-EM map with a resolution of 4 Å. The structure revealed direct interactions between mLST8 and RagA-GTP. Residues His49 and Arg51 of the interswitch region of RagA maintained electrostatic interactions with an acidic patch on mLST8, involving residues Asp181, Asp213, and Glu300 (Extended Data Fig. 5a,b). The ordered interswitch region of RagA in the GTP-bound state ensures that only RagA-GTP can interact with mLST8 (Extended Data Fig. 5c). This novel Rag-Ragulator site partially overlaps the mSIN subunit of mTORC2, consistent with the absence of interaction between Rag-Ragulator and mTORC2 (Extended Data Fig. 6a). This site is near the binding site of the mTORC1 inhibitory protein PRAS40 ^4^ (Extended Data Fig. 6b).

### mTOR and Raptor are membrane-interacting proteins

The cryo-EM reconstruction revealed that mTOR and Raptor subunits directly interact with the membrane, in addition to the expected membrane attachment by lipidated Lamtor1 and Rheb (Fig. 3a). In line with previous publications ^14,15,23–25^, the N-terminal membrane anchoring segment of Lamtor1 was not visualized. The last ordered residue Ala173 of Rheb is in the middle of the hypervariable domain (HVR), indicating a partially folded HVR and a flexible 8-residue loop attached to membrane through Cys181, which places Rheb about 13 Å above the membrane (Fig. 2b). The hydrophobic side-chains Phe1296 and Met1297 in the WD40 domain of Raptor, which we denote as the “FM finger”, manifested membrane interactions in the cryo-EM density map (Fig. 3c). Specific residues in the N-HEAT domain of mTOR, namely Lys471, Arg472, and Lys474, form a basic loop that is also in close contact with the membrane (Fig. 3d). The entire active mTORC1 assembly showed a large membrane-covering footprint, due to the membrane contact sites of mTOR and Raptor subunits, as well as the small membrane-Rheb gap (Fig 3e).

The two Raptor-membrane contact sites in the dimeric structure are separated by 230 Å (Fig 3e). To accommodate both contact sites simultaneously, the liposome diameter must be large enough to provide a relatively flat platform within the potential flexibility range of the complex. Accordingly, we reasoned that liposomes with smaller diameters would result in reduced mTORC1 activation. Liposomes with diameters of 30 or 50 nm showed about a 65% reduction in mTORC1 kinase activity compared to those with a 400 nm diameter (Fig. 3f), bearing out the structural prediction.

To validate the biochemical role of the membrane-interacting residues of Raptor and mTOR, we mutated Raptor residues Phe1296 and Met1297, as well as mTOR residues Lys471, Arg472, and Lys474, to acidic residues in order to maximally disrupt membrane anchoring. We then purified mTORC1 with either the Raptor mutations, the mTOR mutations, or a combination of both, and tested them using an in vitro kinase assay. Mutations in mTORC1 containing Raptor^F1296E/M1297E^ alone, mTOR^K471D/R472D/K474D^ alone, or the combination of both reduced the kinase activity of mTORC1 (Fig. 3g), consistent with the membrane-docked structure. To validate the functional role of the Raptor-membrane interaction, we tested the constructs of Raptor^F1296E^ and Raptor^F1296E/M1297E^ in the inducible Raptor knockout (iRapKO) Mouse Embryonic Fibroblasts (MEFs) (Fig. 3h). After the treatment of 4-hydroxy-tamoxifen (4-OHT), we confirmed the knockout of Raptor. We observed a significant reduction in 4EBP1 and S6K phosphorylation in both Raptor mutants compared to the wild type, upon the replenishment of amino acids. These results collectively show that the full activation of mTORC1 requires the interactions between membrane and mTOR and Raptor subunits.

### Conformational pathway for Rheb activation of mTORC1 on the membrane

Using symmetry expansion and particle subtraction to remove Raptor-Rag-Ragulator, we performed 3D classification on the subpopulation that does not contain extra Rag-Ragulator and identified two distinct conformations of the mTOR-Rheb-mLST8 subcomplex. We designate these states as “intermediate” and “active” for reasons described below. The final resolutions of the intermediate and active states were 3.61 Å and 3.16 Å, respectively (Extended Data Fig. 1).

By aligning these states based on the Rheb structure, we revealed the conformational changes between the intermediate and active states. The M-heat domain of the active state moved more toward the N-heat domain than in the intermediate state, with an average distance of about 5 Å (Fig. 4a; Extended Data Fig. 7a). The FAT and kinase domains of the active state moved about 5-7 Å more toward the Rheb than in the intermediate state. As a result, the mLST8 subunit moved more than 10 Å toward the Rheb in the active state (Extended Data Fig. 7a). The soluble mTORC1-Rheb structure is between the conformational pathway of the intermediate and the active states, when superimposed on the Rheb subunit (Extended Data Fig. 7b).

**Fig. 4:**
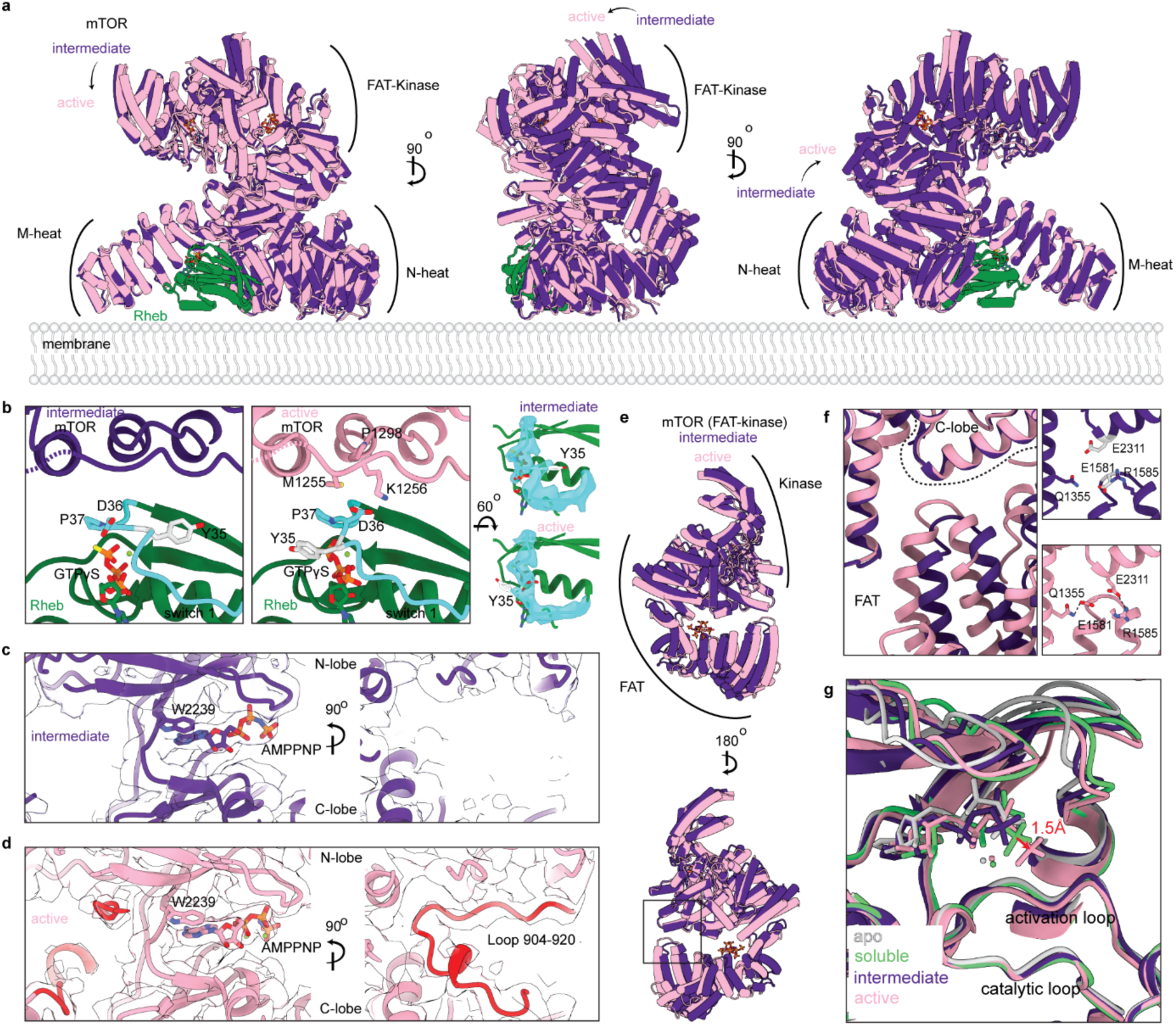
Conformational flexibility of mTOR on membrane. **a**, The intermediate and active states of mTOR are superimposed based on the Rheb subunits. The movement between the intermediate and active states is indicated with arrows from different views. **b**, Close-up view of the interaction between residues 1255-1260 of mTOR and switch I of Rheb in the intermediate and active sites. The cryo-EM density corresponding to switch I is shown. Close-up view of the active site in the intermediate (**c**) and active (**d**) states, the ordered loop (residues 904-920) is colored in red. **e**, The FAT (residues 1255-1453 are omitted) and kinase domains of the intermediate and active states are superimposed based on the C-lobe (residues 2200-2400). **f**, A close-up view of the interactions between FAT and C-lobe domains of mTOR, with insets indicating the residues in intermediate and active states. **g**, Different states of mTOR structures are superimposed based on the C-lobe (residues 2200-2400). The distance of the γ-phosphate of ATP between the soluble mTORC1-Rheb and the active state is shown in red.

The mTOR-Rheb interaction is largely maintained in both intermediate and active states, however, the residues Met1255 and Lys1256 of the FAT domain of mTOR are disordered in the intermediate state but engaged with Rheb in the active state (Fig. 4b). Notably, in the mTOR intermediate state, the residue Tyr35 in the switch I region of Rheb flips to the opposite side and disengaged with the nucleotide, while in the mTOR active state, the switch I region of Rheb adopts the canonical conformation of the Rheb GTP-bound state. In addition, we observed an ordered loop in the active state, located in the opposite site of ATP-binding pocket, while the loop is disordered in the intermediate state and in the soluble mTORC1-Rheb structure (Fig. 4c,d; Extended Data Fig. 7c).

To reveal local conformational differences in the kinase domain and in the ATP-binding pocket, we superimposed the intermediate and active states of mTOR based on the C-lobe of the kinase domain. The FAT domain of mTOR is more engaged with the C-lobe in the active state than in the intermediate state (Fig. 4e; Extended Data Fig. 8a). The established electrostatic interactions between residues Glu1581 and Gln1355 in the FAT domain, as well as between the residues Arg1585 of the FAT domain and Glu2311 of the C-lobe in the active state, indicate direct communication between the FAT domain and the kinase domain for promoting the active state (Fig. 4f). In the ATP-binding pocket, the intermediate state showed similar conformation as the apo state in the ATP-binding pocket, despite the drastic movement of the M-heat domain upon Rheb binding (Fig. 4g). In the active state, the ATP molecule was positioned 1.5 Å closer to the catalytic residues in the C-lobe compared to the soluble mTORC1-Rheb structure, aligning more closely with other atypical serine/threonine kinases in the PIKK family in their active states (Fig. 4g; Extended Data Fig. 8b).

To characterize the relationships between different mTORC1 states and the membrane engagement, we performed three-dimensional variability analysis (3DVA) using the same symmetry-expanded and Raptor-Rag-Ragulator subtracted particles that were used for 3D classification (Extended Data Fig. 9a). We identified one component that does not involve protein stretching into the membrane. We then extracted particles that are at the two ends of this variability component for further local refinement (Extended Data Fig. 9b). The two cryo-EM maps resembled the intermediate and active states identified in the 3D classification, as the absence or presence of the corresponding loop 904-920 is confirmed, respectively (Extended Data Fig. 9c). The active state from 3DVA showed clear density for the basic loop of Lys471, Arg472, and Lys474 and an extended helix, indicating stronger membrane engagement, while the basic loop and the helix are not observed in the intermediate state (Extended Data Fig. 9d). However, the Raptor-membrane interaction is present in both intermediate and active states (Extended Data Fig. 9e). This implies that direct Raptor-membrane interaction is a general requirement for mTORC1 membrane engagement, while the mTOR-membrane interaction promotes the activation of the mTOR kinase.

## Discussion

Among the questions we set out to address was why Rheb-GTP, which is critical for GF-dependent activation of mTORC1 in cells ^3,5^, is such a low-affinity activator in solution ^4^. We found that the physiological concentrations of lipidated Rheb-GTP, in the presence of liposomes, membrane-tethered Ragulator, and an active Rag dimer, could potently activate mTORC1 in a biochemical reconstitution. This could be explained, at least in part, simply by the increased local concentration of Rheb-GTP in the vicinity of mTORC1. Reduction of dimensionality, which limits the search space accessible by diffusion to two dimensions ^26^, could be a contributing factor, but the limitations of this mechanism in biology have been noted ^27^. We probed the mechanism more deeply, however, using cryo-EM structure determination of the reconstituted system on the membrane. We found that precise structural responses to the membrane context are also important.

By carrying out an atomically detailed analysis of more and less active conformations in the membrane context, we were able to map an activation pathway mediated by the membrane itself. We found that both the mTOR and Raptor subunits of mTORC1 directly engage with the membrane. Full membrane engagement by Raptor and mTOR contacts separated by over 230 Å promotes large-scale conformational rearrangements of the N- and M-HEAT, FAT, and kinase domains of mTOR. These large-scale changes in turn re-orient the N- and C-lobes of the kinase domain to fine-tune the kinase active site in a catalytically optimal geometry. The contributions of membrane anchoring and membrane shape to catalytic activation were verified by reconstitution of membrane binding site mutants of Raptor and mTOR and analyzing the shape dependence of liposome activation. Lysosomes undergo tubulation and swelling in the course of their normal function and under stress ^28,29^, which raises the possibility that membrane shape changes could influence mTORC1 activation.

Transduction of the activating membrane engagement signal from the plane of the membrane involves pivoting of the side-chain of Rheb Tyr35. In the intermediate state, the side-chain of Tyr35 is in an unexpected conformation, which points towards mTOR instead of the Rheb GTP binding site. This serves to restrict the close approach of mTOR to Rheb and so to the membrane. The Tyr35 side-chain rotates away from mTOR upon activation, relieving this steric barrier to close engagement. The mutation *RHEB* Y35N was discovered in endometrial and kidney clear cell tumors in the course of a large-scale genetic analysis ^30^. Expression of Rheb Y35N hyperactivates mTORC1 ^31,32^. The structure of Rheb Y35N resembles that of wild-type and the mutant activates mTORC1 in solution to a similar extent ^33^, thus the mechanism of mTORC1 hyperactivation by this mutant has been unknown. Our observations of two activity states of membrane-bound mTORC1 suggest that Rheb Y35N acts by promoting the fully membrane-engaged active conformation of mTORC1.

mTOR is a member of the phosphatidylinositol-3 kinase-related kinase (PIKK) superfamily. While the role of membrane anchors with large spatial separations in kinase activation is unique, similar local rearrangements of the N- and C-lobes of other PIKK family members, ATM, Mec1^ATR^, and DNA-PK, have been reported in response to ROS-dependent activation ^34^, constitutive activating mutation (F2244L) ^35^, and DNA-activation ^36^, respectively. We observed that the ATP molecule in the active state of membrane-bound mTORC1 aligns well with other activated PIKK family members, indicating a more active conformation of mTORC1.

The observation in the membrane-bound structure of a second, novel Rag-Ragulator binding site on mLST8 was unexpected. The mLST8 subunit is common to both mTORC1 and mTORC2, but the additional presence of SIN1 in mTORC2 blocks the novel Rag-Ragulator site, consistent with the known lack of interaction of mTORC2 with Rags. The novel Rag-Ragulator site overlaps with the binding site for proline-rich Akt substrate of 40 kilodaltons (PRAS40) on mLST8 in mTORC1 ^4^. PRAS40 antagonizes GF-dependent mTORC1 activation ^37–40^. It remains to be determined, however, whether the novel Rag-Ragulator site has a physiological activating role, whether via antagonizing PRAS40 inhibition, or simply augmenting lysosomal recruitment. Another open question concerns the role of membrane contacts in mTORC1 phosphorylation of TFEB and other non-canonical substrates, which requires Rag-Ragulator but not Rheb ^6,20,41^.

mTORC1 is activated on lysosomes in response to GF signaling, despite that only a small fraction of the key mediator of GF signaling, Rheb, is transiently localized on lysosomes ^42^ ^43^. Physiologically, this serves to AND-gate the signal to proliferate with the availability of amino acids needed as building blocks for growth. Yet it has been unclear how signal integration is executed at the structural levels. The Rag GTPase dimer is tethered to the lysosome by the Ragulator complex. Ragulator is anchored by lipidation of its Lamtor1 subunit, and the lipid anchor is connected to the folded core of Ragulator by a 45-residue N-terminal disordered region ^23,24,44–46^, with an estimated end-to-end length of ∼47 Å on average and an extended length of ∼100 Å (Extended Data Fig. 10a) ^47^. The distance is potentially longer because of the volume-exclusion effects due to its tethering to the membrane surface ^48^. Augmented to the dimensions of the Rag-Ragulator complex leads us to estimate that the membrane-docking site of mTOR would be tethered within ∼40 Å of the membrane surface upon Rag-Ragulator engagement. This is close enough to dramatically increase the probability of encountering membrane-tethered Rheb, which itself is tethered at a mean distance of ∼15 Å and an extended distance of ∼40 Å from the membrane (Extended Data Fig. 10b). Even Rheb is still not close enough to drive full membrane engagement and activation on its own. Thus, lysosomal membrane engagement is a three-step process, in which initial localization to within a ∼10 nm vicinity of the membrane is driven by Rag-Ragulator. This allows capture by Rheb at a distance of ∼1.5-4 nm from the membrane. The final docking to the membrane is driven by the Raptor and mTOR membrane docking sites, leading to full activation (Fig. 5). This model provides a satisfying structural explanation for how mTORC1 integrates growth factor and nutrient signals in the context of the lysosomal membrane.

**Fig. 5:**
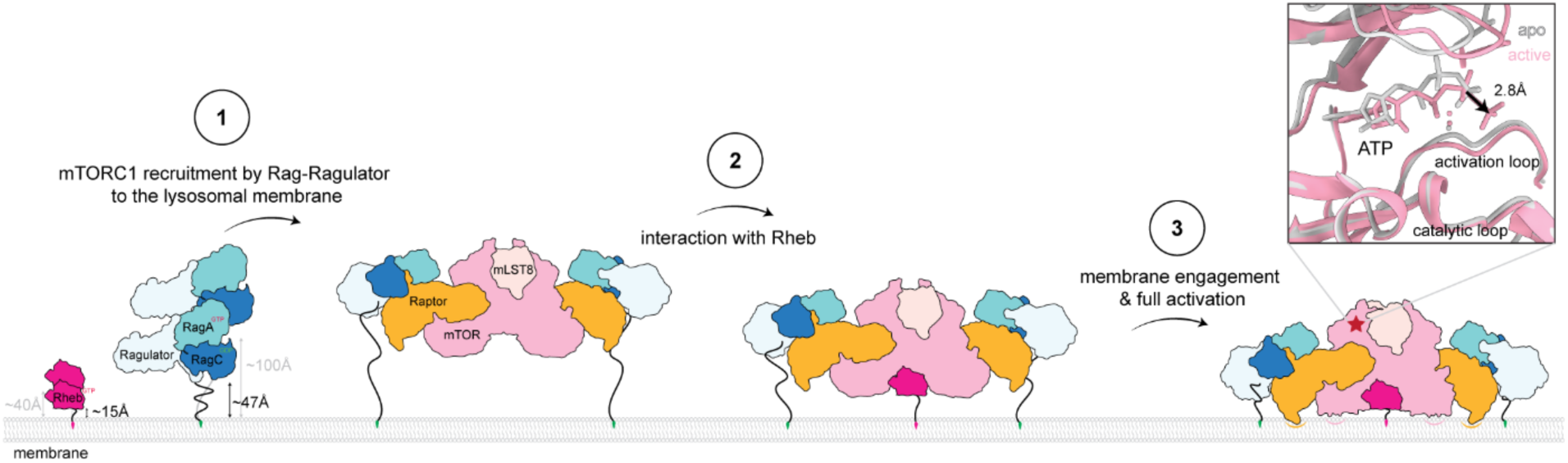
A model of mTORC1 recruitment and activation on the lysosomal membrane. The average distances between the protein and membrane are indicated by black arrows. The linkers that anchor Lamtor1 and Rheb to membranes are arbitrary. The gray arrows indicate the most possible extended positions relative to the membrane surface. The insect shows the ATP binding site of mTORC1 in the apo and active states.

## Acknowledgements

We thank D. Toso and R. Thakkar for cryo-EM facility support, and members of the Hurley laboratory for insightful discussions. This work was supported by Genentech as part of the Alliance for Therapies in Neuroscience and the National Cancer Institute NIH, R01 CA285366 (J.H.H.); the Italian Telethon Foundation (to G.N., and A.B.); Fondazione AIRC per la ricerca sul cancro (MFAG-23538 to G.N.; IG-29230 to A.B.); MIUR (PRIN 2022CRFNCP to G.N.; PRIN 202032AZT3 to A.B.; PRIN P2022T4PKT to G.N. and A.B.); Wereld Kanker Onderzoek Fonds (WKOF), as part of the World Cancer Research Fund International grant program (IIG_FULL_2022_009 to G.N.); and the European Research Council (INCANTAR - Grant # 101097752 to A.B.).

## Data availability

Structural coordinates will be deposited in the RCSB and EM density maps in the EMDB prior to publication. (Deposition will start soon.)

## Methods

### Protein expression and purification

The full-length, codon-optimized genes of human RagC with the S75N mutation and human RagA with the Q66L mutation were synthesized (Twist Bioscience) and individually cloned into a pCAG vector. Mutations were introduced in RagA (Q66L) and RagC (S75N) to lock the active state of Rag GTPases (RagA^GTP^–RagC^GDP^). The RagC (S75N) construct included a TEV-cleavable GST tag at the N terminus, while the RagA (Q66L) was tagless. For the expression and purification of the Rag GTPases, HEK293F GnTI-cells were transfected with a total of 1 mg plasmid DNA (550 μg RagA, and 450 μg RagC) and 4 mg polyethylenimine (PEI) (Sigma-Aldrich) per liter at a density of 1.8 × 10^6 cells/ml. Cells were collected after 72 hours and lysed via gentle nutation in wash buffer (50 mM HEPES, 150 mM NaCl, 2.5 mM MgCl_2_, 1 mM TCEP, pH 7.4) supplemented with 0.4% CHAPS and Protease Inhibitor (Roche) for 1 hour. The lysate was cleared by centrifugation at 35,000 × g for 35 minutes. The supernatant was incubated with glutathione Sepharose 4B (GE Healthcare) resin for 2 hours. The resin was then washed first with a modified wash buffer (200 mM NaCl and 0.3% CHAPS) and then with the wash buffer. The complex was eluted by on-column TEV-cleavage overnight without nutation. Eluted complexes were concentrated and further purified by size exclusion chromatography (SEC) using a Superose 200 10/300 GL (GE Healthcare) column equilibrated with the wash buffer.

The full-length, codon-optimized genes of the human Ragulator complex (Lamtor1-5) were synthesized (Twist Bioscience) and individually cloned into a pCAG vector. The Lamtor1 construct includes a TEV-cleavable GST tag at the N terminus, followed by a His6 tag, which replaces its first seven residues. The Lamtor2 construct features a TEV-cleavable tandem 2× Strep II-1× FLAG-tag (2SF-TEV) at the N terminus. A total of 1 mg plasmid DNA (200 μg each of the five constructs) and 4 mg polyethylenimine (PEI) (Sigma-Aldrich) were used to transfect HEK293F GnTI-cells at a density of 1.8 × 10^6 cells/ml per liter. The cells were pelleted after 72 hours of transfection and lysed in wash buffer containing 1% Triton X-100 and protease inhibitor. The cleared supernatant after centrifugation was applied to glutathione Sepharose 4B (GE Healthcare) resin and incubated for 2 hours. The complex was eluted by on-column TEV cleavage overnight without nutation and supplemented with a final concentration of 0.5 mM EDTA. Further purification was performed by SEC using a Superdex 200 10/300 GL column equilibrated with wash buffer. The fractions containing all five subunits were pooled and concentrated.

To assemble the Rag-Ragulator complex, an excess amount of Rag was incubated with Ragulator and Ni-nitrilotriacetic acid resin (Thermo Scientific) at 4°C for 1 hour. Excess Rag was removed by washing the resin with wash buffer containing 40 mM imidazole. The assembled Rag-Ragulator complex was eluted using wash buffer with 250 mM imidazole and further purified by SEC with a Superdex 200 10/300 GL column.

The human mTORC1 complex was purified similarly to previous methods ^20^. The full-length, codon-optimized genes for human mTOR, Raptor, and mLST8 were cloned into pCAG vectors (mTOR with TEV-cleavable 2SF, Raptor with uncleavable 2SF, and mLST8 with uncleavable 2SF). A total of 1.35 mg plasmid DNA (900 μg mTOR, 250 μg Raptor, and 200 μg mLST8) and 4 mg polyethylenimine (PEI) (Sigma-Aldrich) were used to transfect HEK293F GnTI-cells at a density of 1.8 × 10^6 cells/ml per liter. Cells were harvested 72 hours post-transfection and lysed in the same buffer used for RagAC purification. The complex was purified using Strep-Tactin resin (IBA Lifesciences) and eluted with wash buffer containing 10 mM D-desthiobiotin. The elution was then diluted into an equal volume of salt-free buffer (50 mM HEPES, 1 mM TCEP, pH 7.4) and applied to a 1 ml HiTrap Q column (GE Healthcare). The mTORC1 complex and free RAPTOR were separated by a 20 ml salt gradient to a final concentration of 0.5 M NaCl, using the salt-free buffer and high-salt buffer (50 mM HEPES, 1 M NaCl, 1 mM TCEP, pH 7.4). The flow rate was 0.2 ml/min, and the fraction size was 0.2 ml. The fractions containing the mTORC1 complex and free Raptor were collected and concentrated with Amicon^®^ Ultra-4 concentrators. Mutations in the mTOR and Raptor genes were generated using NEBuilder^®^ HiFi DNA assembly. The mTORC1 complex mutants were produced by altering the combination of wild-type and mutated genes during transfection and purified as described above.

The plasmids containing full-length human 4EBP1 and Rheb genes were gifts from the Zoncu lab. These genes were individually cloned into a 2GT vector from QB3 Macrolab (https://qb3.berkeley.edu/facility/qb3-macrolab/), which features a TEV-cleavable tandem GST-His6 tag at the N-terminus. 4EBP1 and Rheb were overexpressed in *Escherichia coli* Rosetta 2(DE3) strains and purified using the same method. The *E. coli* cells were grown in LB medium at 37°C until an OD of 0.6, then induced by adding 0.2 mM IPTG at 18°C overnight. Cells were collected, resuspended in Ni buffer (50 mM Tris-Cl pH 7.5, 300 mM NaCl, 20 mM imidazole, 5 mM 2-mercaptoethanol, 1 mM PMSF), and lysed by sonication. The protein was purified using a HisPur Ni-NTA Resin (Thermo Scientific), washed with Ni buffer containing 40 mM imidazole, and eluted with 250 mM imidazole. The GST-His6 tag was cleaved by incubating with TEV enzyme overnight. Further purification was performed by SEC using a Superdex 75 10/300 GL column equilibrated with buffer (50 mM HEPES, pH 7.5, 150 mM NaCl, 0.5 mM TCEP). Fractions containing the desired proteins were passed through the Ni column to remove residual GST-His6 and then concentrated.

To charge Rheb with GTPγS (Abcam) or GDP (Sigma-Aldrich), Rheb was first diluted in buffer (50 mM HEPES, pH 7.5, 150 mM NaCl, 1 mM TCEP, 5 mM EDTA) and then supplemented with GDP or GTPγS at a 30-fold molar excess. The mixture was incubated at 30°C for 1 hour, followed by the addition of 20 mM MgCl_2_. Further purification was done by SEC using a Superose 75 10/300 GL (GE Healthcare) column to remove excess nucleotides.

All steps of protein purification were performed at 4°C and aliquoted proteins were flash frozen in liquid nitrogen and stored at −80°C.

### Large unilamellar vesicles (LUVs) preparation

A lipid mixture was prepared in a glass vial using the lipid composition shown in Extended Data Table 1. To form a thin film on the glass wall, the glass vial was slowly shaken on a vortex while drying under nitrogen gas. The glass vial was then placed in a vacuum oven overnight at room temperature to evaporate any remaining solvent. The lipids were hydrated in a lipid buffer (25 mM HEPES pH 7.2, 100 mM NaCl) to a final concentration of 1.8 mM for 1 hour, with intermittent vortexing during hydration. The solution was transferred to a 15 mL Eppendorf tube and subjected to 9 cycles of freeze/thaw using liquid nitrogen and a 40°C water bath. The lipid mixture was either stored at −80°C or immediately extruded using an Avanti Polar Lipids Mini Extruder (Cat No. 610023) at least 40 times through a 200 nm filter (Whatman® NucleporeTM Track-Etched Membranes, diam. 19 mm) for mTORC1 kinase activity assays and cryo-EM studies. Filters of different diameters were also used to generate various sizes of liposomes to assess the effect of liposome size on mTORC1 kinase activity. The lipid solution after extrusion was stored at 4°C for up to two weeks.

### mTORC1 kinase activity with LUVs

The kinase assay was conducted in a buffer containing 25 mM HEPES (pH 7.2), 100 mM NaCl, 10 mM imidazole, 10 mM MgCl_2_, and 2 mM DTT, at 30°C for 10 minutes, in a final volume of 50 μL. First, Rheb was incubated with liposomes at concentrations of 0.25 μM and 0.18 mM, respectively, at 4°C overnight in 40 μL lipid buffer. In parallel, Rheb alone and liposomes alone were diluted to the same concentrations in the lipid buffer and incubated at 4°C overnight. Following the overnight incubation, the reactions were stopped with 2 mM DTT at room temperature for 30 minutes, followed by the addition of 10 mM MgCl_2_ and 10 mM imidazole. The His_6_-tagged Rag-Ragulator complex was then added to the reactions and incubated on ice for 30 minutes. Subsequently, 4EBP1 and mTORC1 were added to the reactions at final concentrations of 10 μM and 5 nM, respectively. The reactions were initiated by adding ATP to a final concentration of 1 mM and incubated in a thermocycler (Bio-Rad T100) at 30°C for 10 minutes. The reactions were stopped by diluting 10-fold into a urea-denaturing buffer (50 mM Tris-Cl, pH 7.5, 150 mM NaCl, 8 M urea). All reactions were then diluted into 4x NuPAGE LDS sample buffer and boiled for 2 minutes. The proteins were resolved on a 4–12% NuPAGE Bis-Tris gel and transferred to PVDF membranes using the Trans-Blot Turbo Transfer System. Western blotting was performed using the anti-phospho 4EBP1 antibody (Cell Signaling Technology #2855). For Dotblot analysis, 2 μL of the denatured sample in urea buffer was directly applied to nitrocellulose membranes and detected with the same 4EBP1 antibody.

### Cryo-EM sample preparation and imaging

The mTORC1–Rheb–Rag–Ragulator–4EBP1 complex on liposomes was reconstituted in the following steps. First, a mixture of Rheb and liposomes (200 nm) was incubated at 4°C overnight in lipid buffer at concentrations of 8 μM and 1.8 mM, respectively. The Rheb-liposome mixture was supplemented with 2 mM DTT and 10 mM MgCl_2_ and incubated at room temperature for 30 minutes. In parallel, mTORC1 was incubated with His_6_-Rag-Ragulator in lipid buffer containing 5 mM TCEP, at concentrations of 1 μM and 4 μM, respectively. Equal volumes of the Rheb-liposome mixture and the mTORC1-Rag-Ragulator mixture were then combined and incubated on ice for 2 hours. Finally, 4 μM of 4EBP1 and 1 mM of AMPPNP were added to the mixture for 10 minutes before applying it to cryo-EM grids for vitrification.

Cryo-EM samples were prepared by applying 3 μL of the aforementioned complex to a glow-discharged (PELCO easiGlow, 45 s in air at 15 mA and 0.37 mbar) holey carbon grid (C-flat, 2/1-3C-T) and vitrified using an FEI Vitrobot Mark IV (Thermo Fisher Scientific). The samples were incubated on grids for 1 minute and blotted for 3 seconds with blot force 15, using two Whatman 595 papers on the sample side and one Whatman 595 paper on the backside, at 6°C with 95% relative humidity.

Cryo-EM images of the mTORC1–Rheb–Rag–Ragulator–4EBP1 complex on liposomes were recorded using a Titan Krios G3 microscope (Thermo Fisher Scientific) equipped with a Gatan Quantum energy filter (slit width 20 eV) and operated at 300 kV. Automated data acquisition was performed using SerialEM ^49^ on a K3 Summit direct detection camera (Gatan) in super-resolution correlated-double sampling mode with a pixel size of 0.52 Å and a defocus range of −0.9 to −2.2 μm. A total of 36 exposures per stage shift were enabled by large beam shift. The beam intensity was adjusted to a dose rate of approximately 1 e^−^ per Å^^2^ per frame for a 30-frame movie stack with a total exposure time of 5.4 seconds. A total of 58,092 movies were recorded.

### Cryo-EM data processing

The data processing scheme for the mTORC1–Rheb–Rag–Ragulator-4EBP1 complex on liposomes using cryoSPARC v.4 ^50^ is shown in Extended Data Fig 1 and statistics are summarized in Extended Data Table 2. Due to the uneven distribution of liposomes within grid squares, 29,301 micrographs containing liposomes were manually selected for processing. Owing to the dataset’s size, micrographs were split and processed following the same protocol, then combined at the homogeneous refinement stage. Initially, the Blob picker was used to maximize the number of particles. Two-dimensional (2D) classification was employed only to remove obvious ‘junk’ particles (e.g., ice and chaperonin contaminants). An initial model of mTORC1 from an ab initio reconstruction in a previous dataset and three bad initial models from ab initio reconstruction in this dataset were used for heterogeneous refinement. Iterative heterogeneous refinement was performed to select good particles, retaining rare views that may not be identified in 2D classification. To minimize the membrane density from biasing the alignment in the first iterations of refinement, the original initial models were used instead of reconstructions from each heterogeneous refinement. After extensive cleaning using heterogeneous refinement, particles were merged, and duplicates were removed with a 50-Å cutoff distance. Additional heterogeneous refinement was used to further sort out particles. Homogeneous refinement was then performed for the full dataset. After identifying a good particle set, it was used to train TOPAZ particle picking ^51^. Particles from TOPAZ picking underwent the same sorting procedure and were merged with the blob-picked particles, with duplicates removed using a 50-Å cutoff distance.

In total, 337,347 particles containing protein complexes and membranes were selected, and reference-based motion correction was used to produce polished particles ^52^. Symmetry expansion, particle subtraction, and local refinement were used to produce an overall cryo-EM map of an asymmetric unit. Focused 3D classification was then used to separate particles without extra Rag-Ragulator and with either one or two copies of extra Rag-Ragulator. Further particle subtraction and local refinement were used to focus on either mTOR-Rheb-mLST8 or Raptor-Rag-Ragulator subcomplexes. Focused 3D classification was used to separate the intermediate and active states of mTOR. The populations containing extra copies of Rag-Ragulator were pooled together, and further particle subtraction and local refinement were used to obtain the cryo-EM map of mLST8-Rag-Ragulator.

In summary, 179,506 particles were refined to 3.23 Å with a C2 symmetry for the mTORC1-Rheb-Rag-Ragulator-4EBP1 complex without extra Rag-Ragulator on the membrane. Local refinement of the mTOR-Rheb-mLST8 (359,012 particles after symmetry expansion) and Raptor-Rag-Ragulator (189,975 particles after symmetry expansion) yielded maps of 3.12 Å and 2.98 Å, respectively. Further classification of mTOR-Rheb-mLST8 identified the active and intermediate conformations. The final refinement of the mTOR-Rheb-mLST8 in active (133,193 particles after symmetry expansion) and intermediate conformations (109,105 particles after symmetry expansion) results in the resolutions of 3.16 Å and 3.61 Å, respectively. The final resolutions for the mTORC1–Rheb–Rag–Ragulator-4EBP1 complex containing either one (128,189 particles) or two copies (29,652 particles) of extra Rag-Ragulator are 3.81 Å (C2 symmetry) and 3.47 Å (C1 symmetry), respectively. 3D variability analysis for the populations without extra Rag-Ragulator was performed in cryoSPARC v.4 (Extended Data Fig 9).

The overall resolution of all these reconstructed maps was assessed using the gold-standard criterion of Fourier shell correlation ^53^ at a 0.143 cutoff ^54^. Local resolution estimation was done in cryoSPARC v.4.

### Atomic model building and refinement

To build the atomic model for the mTORC1-Rheb–Rag–Ragulator-4EBP1 complex on the membrane, we first fit our previous models into the cryo-EM map as a rigid body using UCSF ChimeraX ^55^, with substituted models of mTOR and Raptor from AlphaFold2 prediction ^56^. A composite map combining the local refinement maps was assembled in UCSF ChimeraX. Model refinement against local maps was performed using PHENIX for real-space refinement ^57^. Manual model building was conducted with COOT ^58^ and ISOLDE ^59^ to iteratively inspect and improve local fitting. All figures and videos were created using UCSF ChimeraX.

### Cell culture

Inducible Raptor KO MEFs were kindly provided by Michael N. Hall (University of Basel, Switzerland). MEFs were cultured in DMEM high glucose (cat. no. ECM0728L, Euroclone) supplemented with 10% inactivated FBS (cat. no. ECS0180L, Euroclone), 2 mM glutamine (cat. no. ECB3000D, Euroclone), penicillin (100 IU/ml) and streptomycin (100 μg/ml) (cat. no. ECB3001D, Euroclone) and maintained at 37 °C and 5% CO_2_. All Raptor mutants used in these cellular assays were generated by using QuikChange II-E Site-Directed Mutagenesis Kit (no. 200555, Agilent Technologies). Cells were transfected in 10cm dishes using using Lipofectamine 2000 transfection reagent (Invitrogen).

### Cell treatment

For experiments involving amino acid starvation, cells were rinsed twice with PBS and incubated for 60 min in amino acid-free DMEM (cat. no. MBS6120661) supplemented with 10% dialysed FBS. Serum was dialysed against 1× PBS through 3,500-molecular weight cut-off dialysis tubing to ensure absence of contaminating amino acids. For amino acid refeeding, cells were re-stimulated for 30 min with 1× water-solubilized mix of essential (cat. no.11130036, Thermo Fisher Scientific) and non-essential (cat. no. 11140035, Thermo Fisher Scientific) amino acids resuspended in amino-acid-free DMEM supplemented with 10% dialysed FBS, plus glutamine.

### Cell lysis, western blotting, and quantification

Cells were rinsed once with PBS and lysed in ice-cold lysis buffer (250 mM NaCl, 1% Triton, 25 mM Hepes pH 7.4) supplemented with protease and phosphatase inhibitors. Total lysates were passed 10 times through a 25-gauge needle with syringe, kept at 4 °C for 10 min and then cleared by centrifugation in a microcentrifuge (14,000 rpm at 4 °C for 10 min). Protein concentration was measured by Bradford assay. Densitometry analysis has been performed to calculate the intensity of phosphorylated and total proteins by using ImageJ software. The ratios between the values of phosphorylated and total proteins were normalized to a control condition. Values of quantitative graphs are mean ± standard error of at least three independent experiments. For statistical analysis, Two-way ANOVA and Dunnett’s post hoc test were used to compare differences between groups that have been split into two factors.

---

**Extended Data Fig.1.**
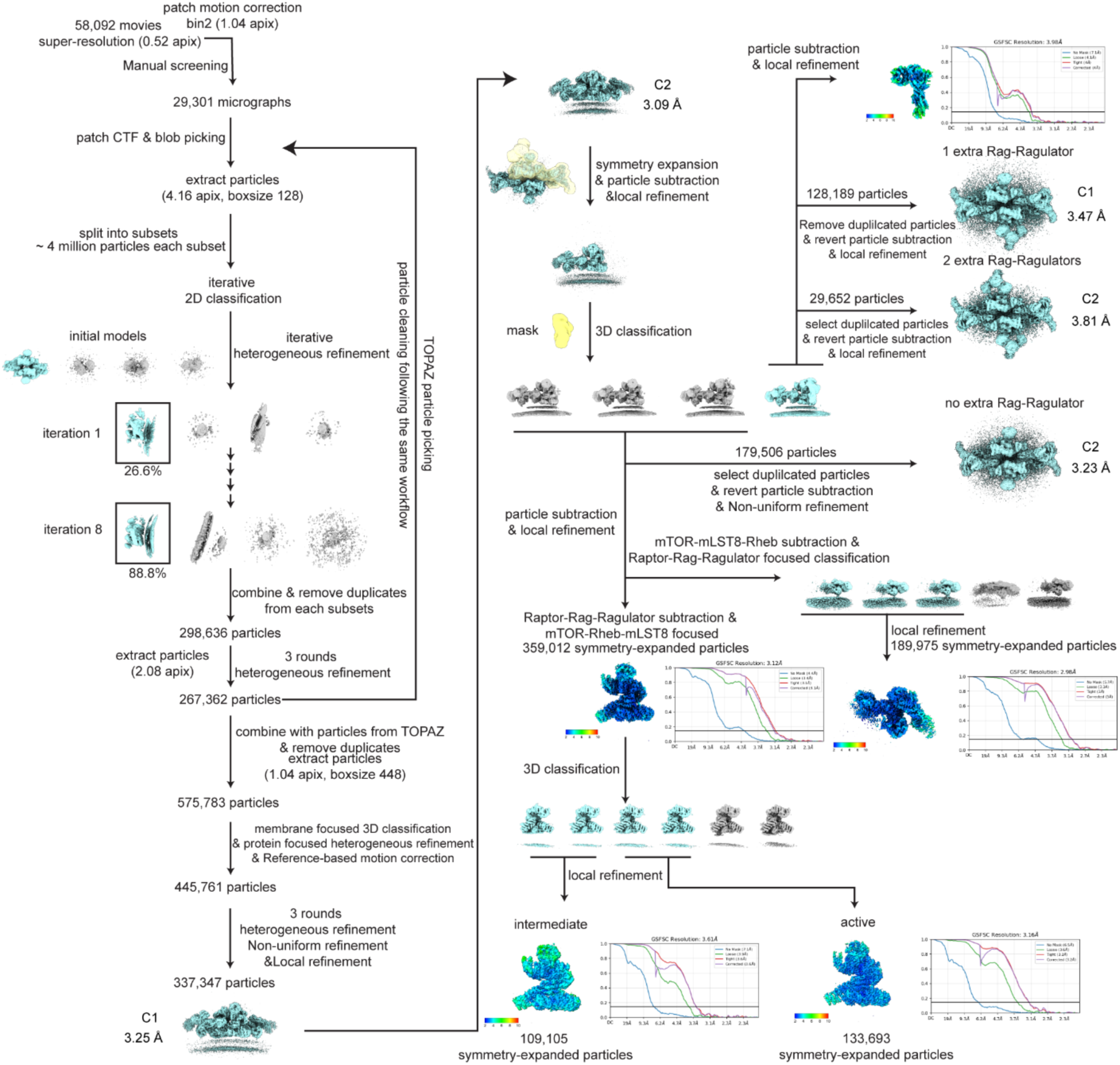
Cryo-EM workflow of mTORC1-Rheb-Rag-Ragulator-4EBP1 on membrane.

**Extended Data Fig. 2.**
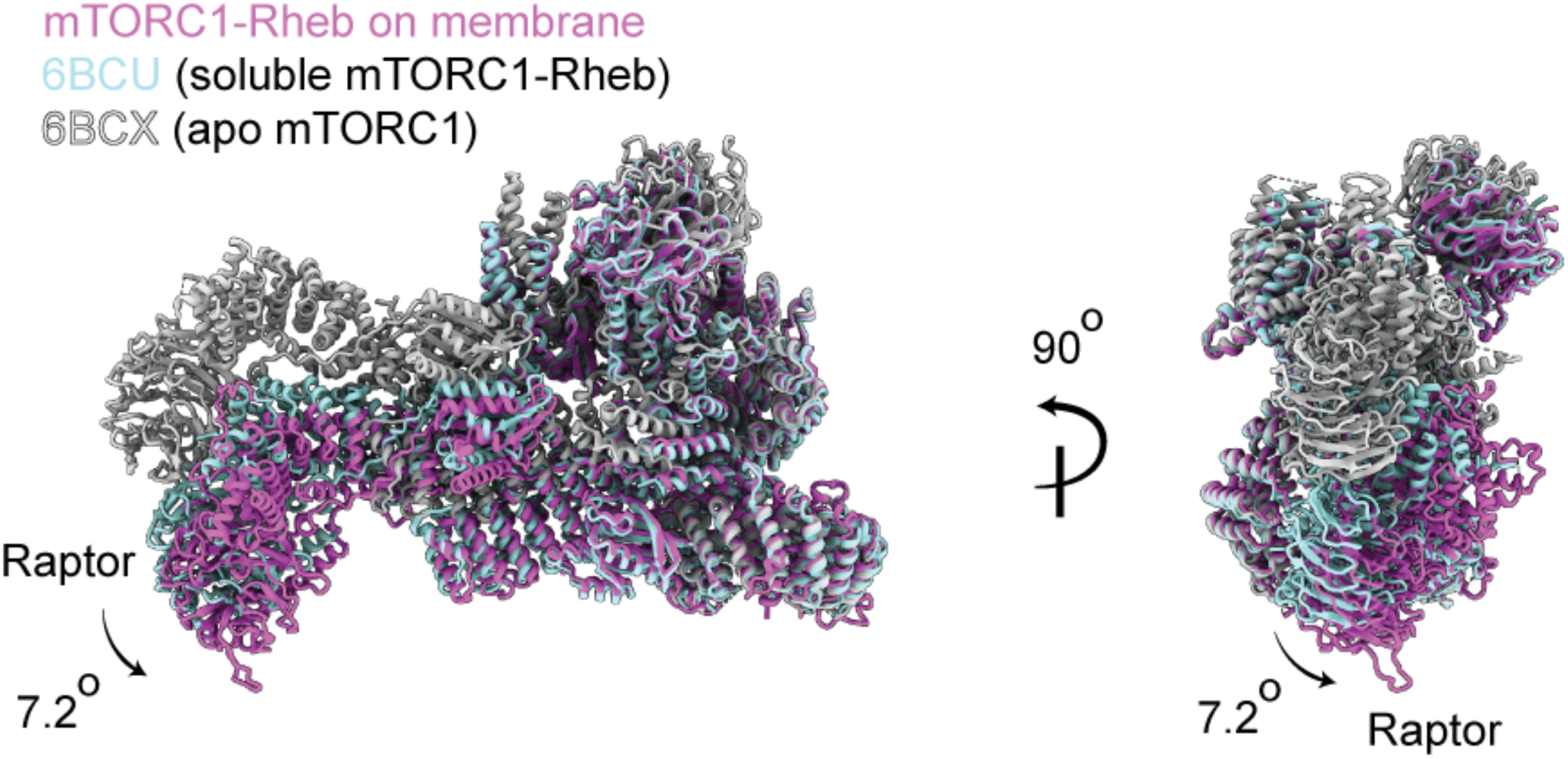
Comparison between mTORC1-Rheb on membrane and in solution. Asymmetric units of apo mTORC1, mTORC1-Rheb in solution, and mTORC1-Rheb on membrane are aligned based on the mTOR subunit. The rotational direction of Raptor subunit between soluble and membrane-bound mTORC1-Rheb is indicated with an arrow.

**Extended Data Fig. 3.**
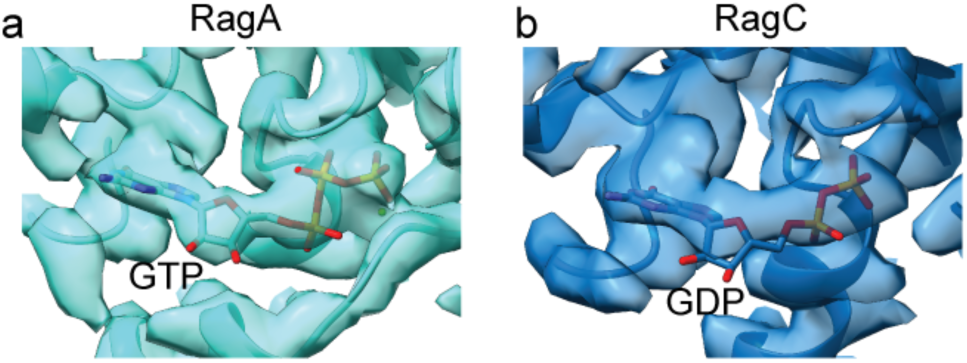
Representative cryo-EM density of the active sites of Rag GTPases. Cryo-EM density of RagA (**a**) and RagC (**b**) at contour level 0.25 and level 0.2, respectively.

**Extended Data Fig. 4.**
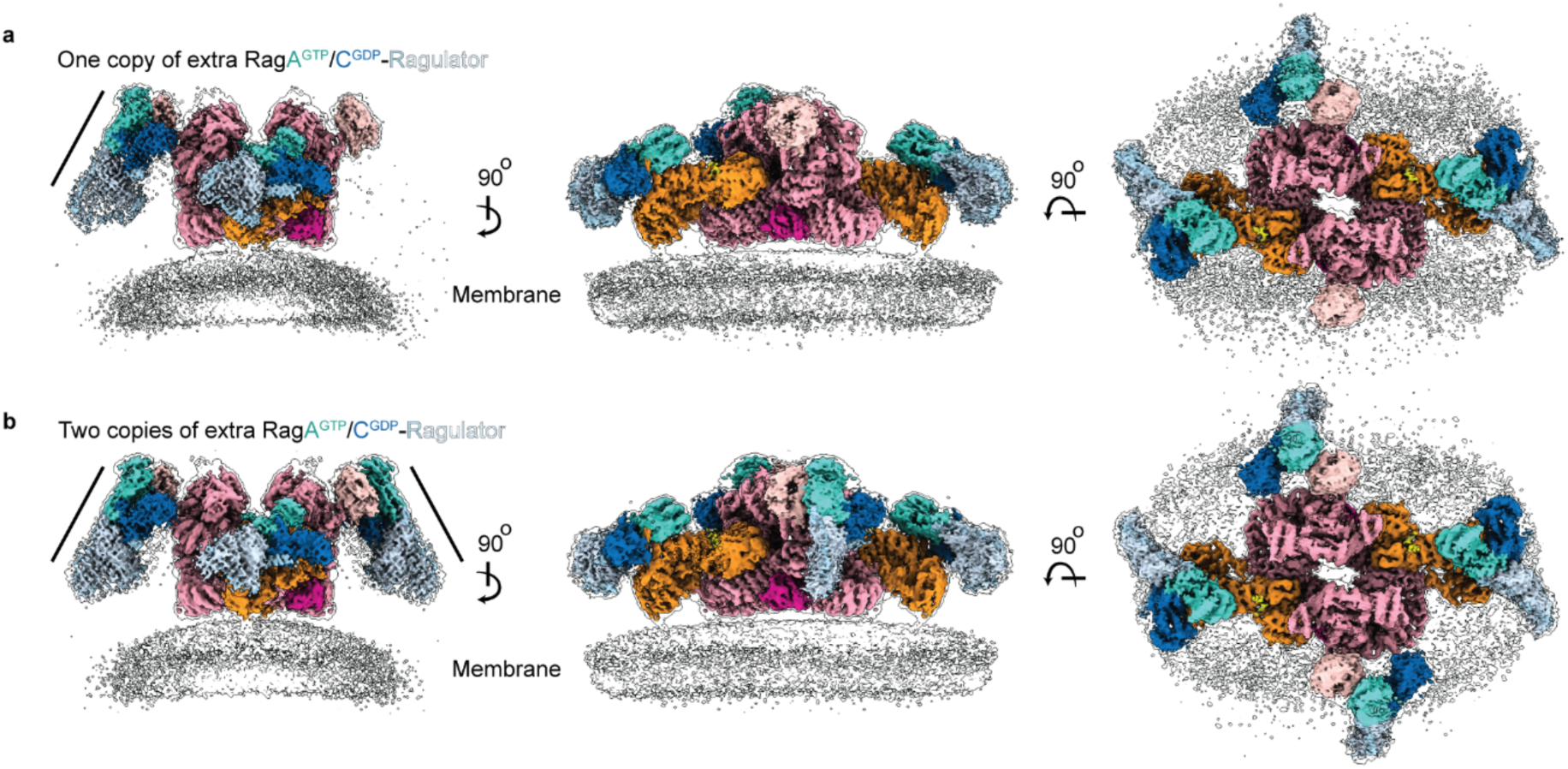
Cryo-EM maps of mTORC1-Rheb-Rag-Ragulator-4EBP1 with extra copy of Rag-Ragulator. **a,** Cryo-EM density map of mTORC1-Rheb-Rag-Ragulator-4EBP1 on membrane with one extra copy of Rag-Ragulator, overlaid with the unsharpened map from the overall refinement with C1 symmetry. **b,** Cryo-EM density map of mTORC1-Rheb-Rag-Ragulator-4EBP1 on membrane with one extra copy of Rag-Ragulator, overlaid with the unsharpened map from the overall refinement with C2 symmetry.

**Extended Data Fig. 5.**
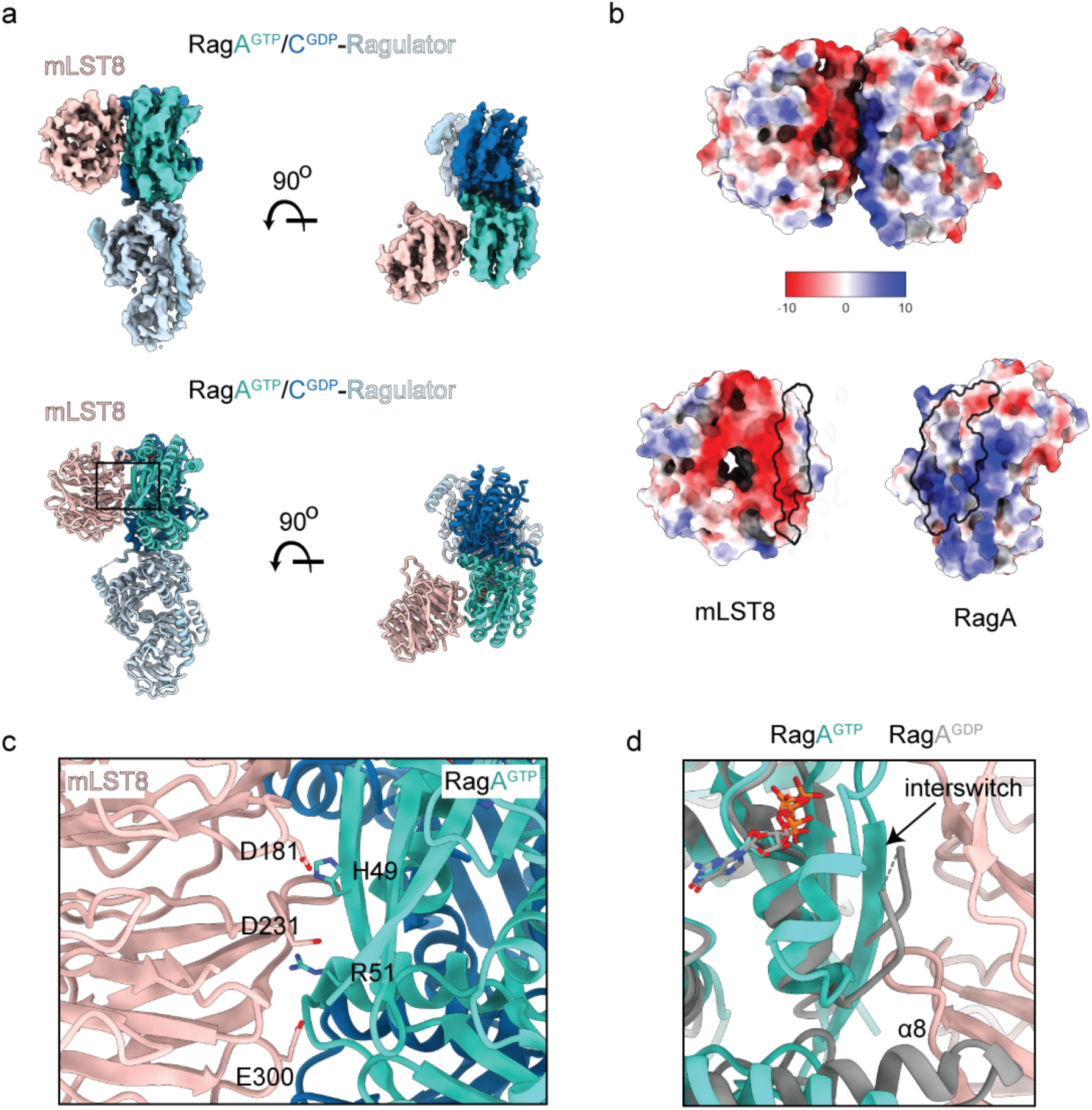
The interaction between mLST8 and RagA. **a,** Cryo-EM density map and model of mLST8-Rag-Ragulator subcomplex. **b,** Electrostatic surface potentials of mLST8 and RagA are shown. The interaction surfaces are outlined. **c,** Close-up view of the interaction between mLST8 and RagA. **d,** Overlay between GTP-bound and GDP-bound RagA (PDB: 6NZD), the missing interswitch region is indicated with arrow. The GDP-bound RagA has a disordered interswitch region, which is responsible for the interaction with mLST8. In addition, the α8 helix in the RagA-GDP has potential clash with mLST8.

**Extended Data Fig. 6.**
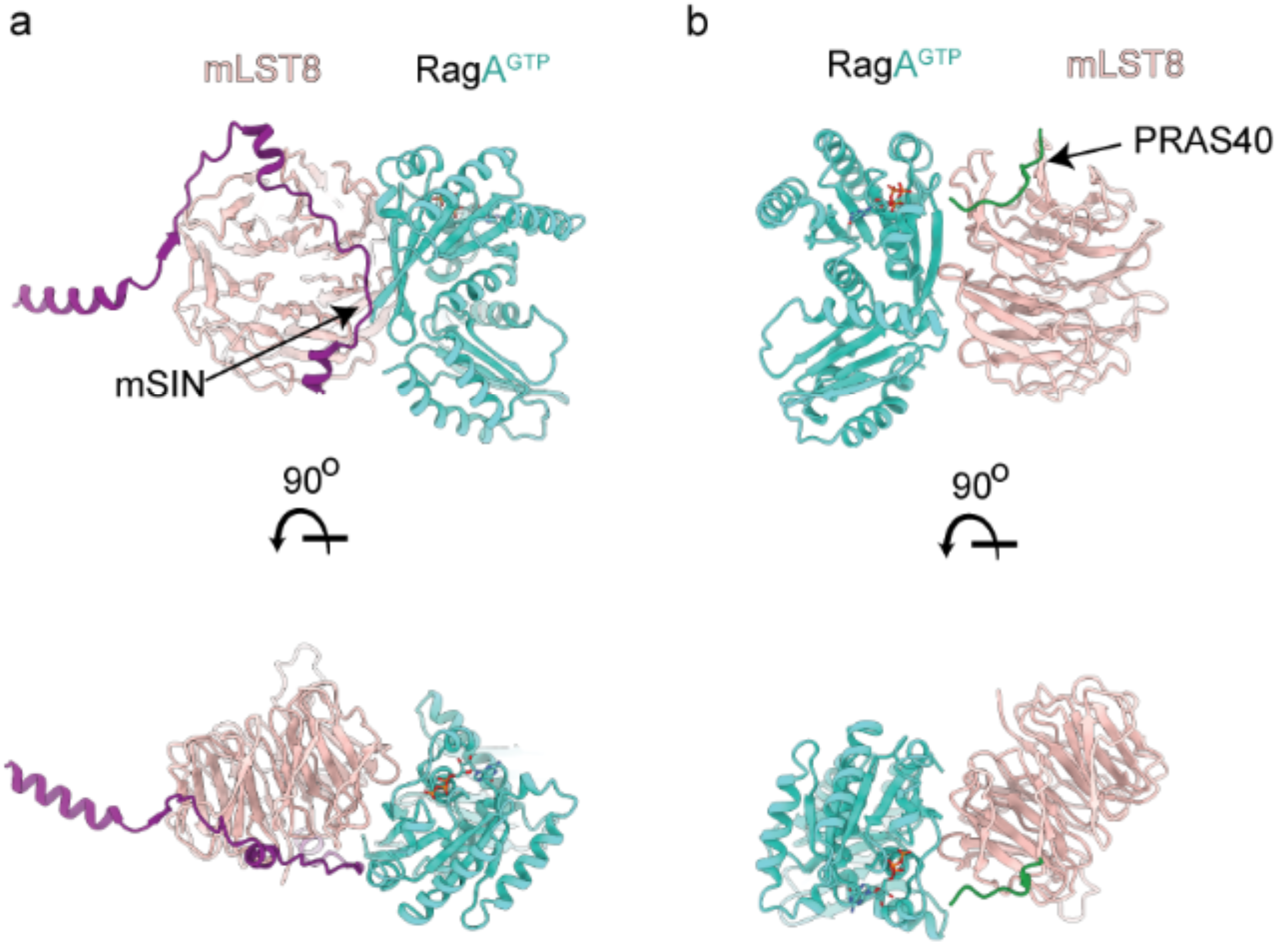
Relative position of mSIN and PRAS40 to mLST8-bound RagA. Structures are superimposed based on mLST8. **a,** A potential clash is observed between mSIN and RagA, indicating this interaction is not compatible with mTORC2. **b,** The mLST8-interacting fragment of PRAS40 is localized close to the mLST8-bound RagA.

**Extended Data Fig. 7.**
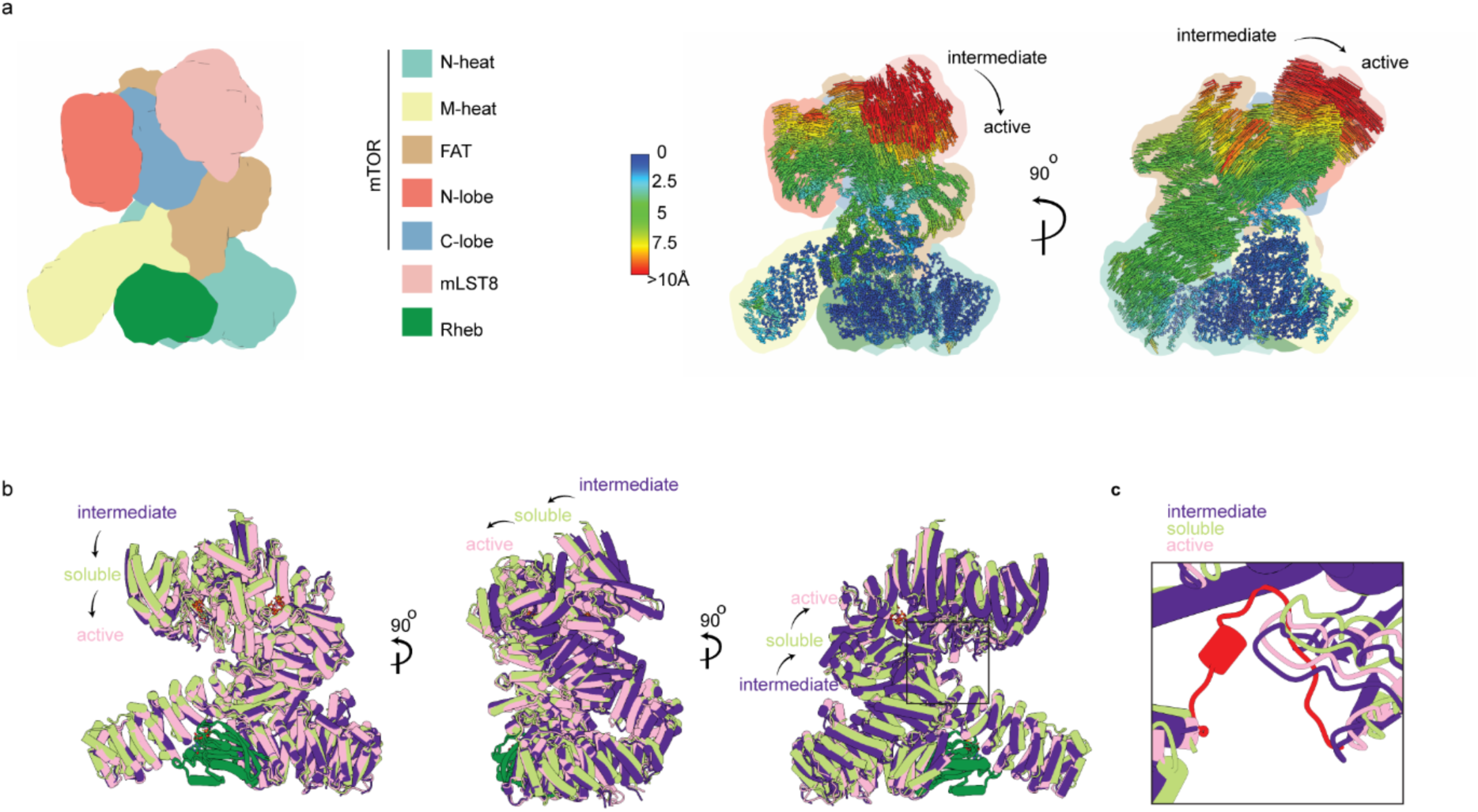
Structural comparison between the intermediate and active states of mTOR. Structures are superimposed based on Rheb. **a,** A carton representation of the mTOR-Rheb-mLST8 subcomplex. The motion between intermediate and active states are indicated with lines that connect the backbone of both structures. The distance of the motion is labeled with a rainbow color. **b,** The soluble structure of mTORC1-Rheb is in between the conformational change of the intermediate and active states of mTORC1-Rheb on membrane. **c,** The soluble structure does not contain the ordered loop in the backside of the ATP pocket. The ordered loop (902-920) is colored red.

**Extended Data Fig. 8.**
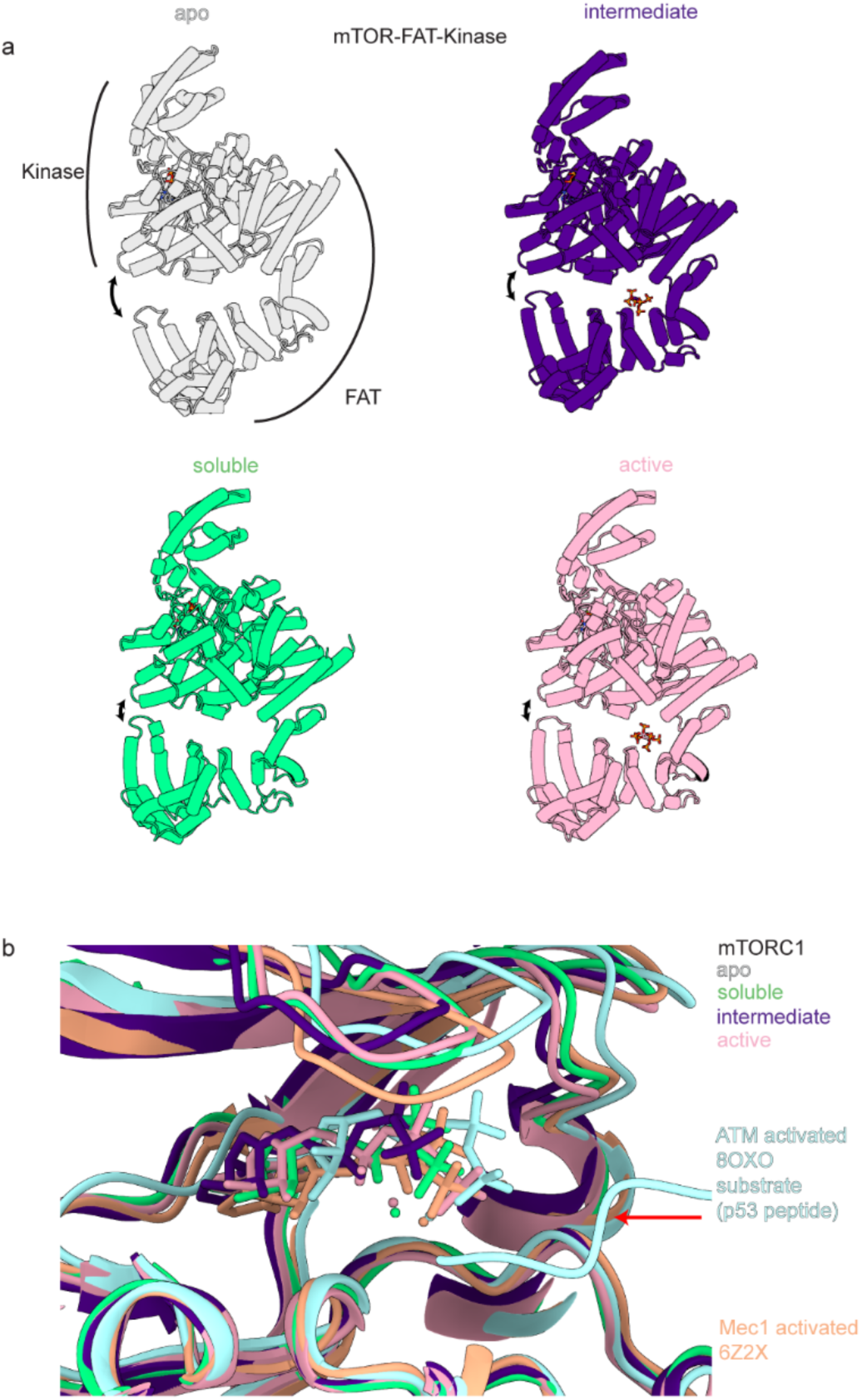
Comparison of the FAT-Kinase domain of mTORC1 in different states and the ATP position relative to other PIKKs. **a,** The FAT (residues 1255-1453 are omitted) and kinase domains of the apo, soluble, intermediate, and active states of mTOR are shown side by side. The constriction between the FAT and kinase domain is indicated by double arrow curved lines. **b,** A close-up view of the ATP binding pocket of mTOR, superimposed by ATM and Mec1 based on the C-lobe (residues 2200-2400). The substrate of ATM, p53, is indicated with a red arrow.

**Extended Data Fig. 9.**
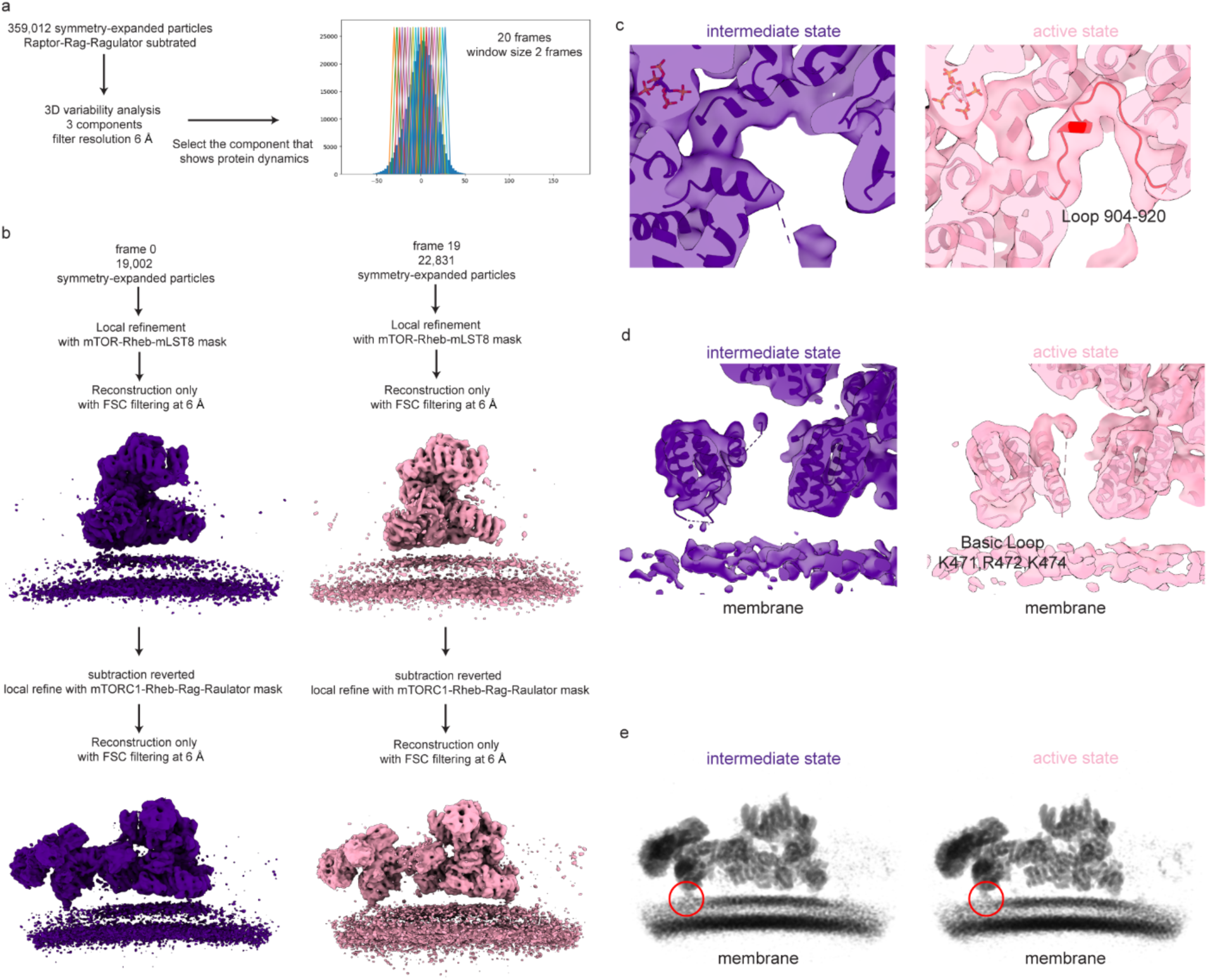
3D variability analysis. **a,** 3DVA processing workflow. **b,** Particles corresponding to the frames at the two ends of the continuous motion were used for local refinement. Final refinement was done with FSC filtering at 6 Å. **c**, Close-up view of the ordered loop region in the opposite of ATP binding pocket. The ordered loop is colored red. **d,** Close-up view of the membrane-interacting site of mTOR. **e,** A slab representation showing the intermediate and active states with Raptor-Rag-Ragulator subcomplex. The membrane-interacting site of Raptor is indicated by red circle.

**Extended Data Fig. 10.**
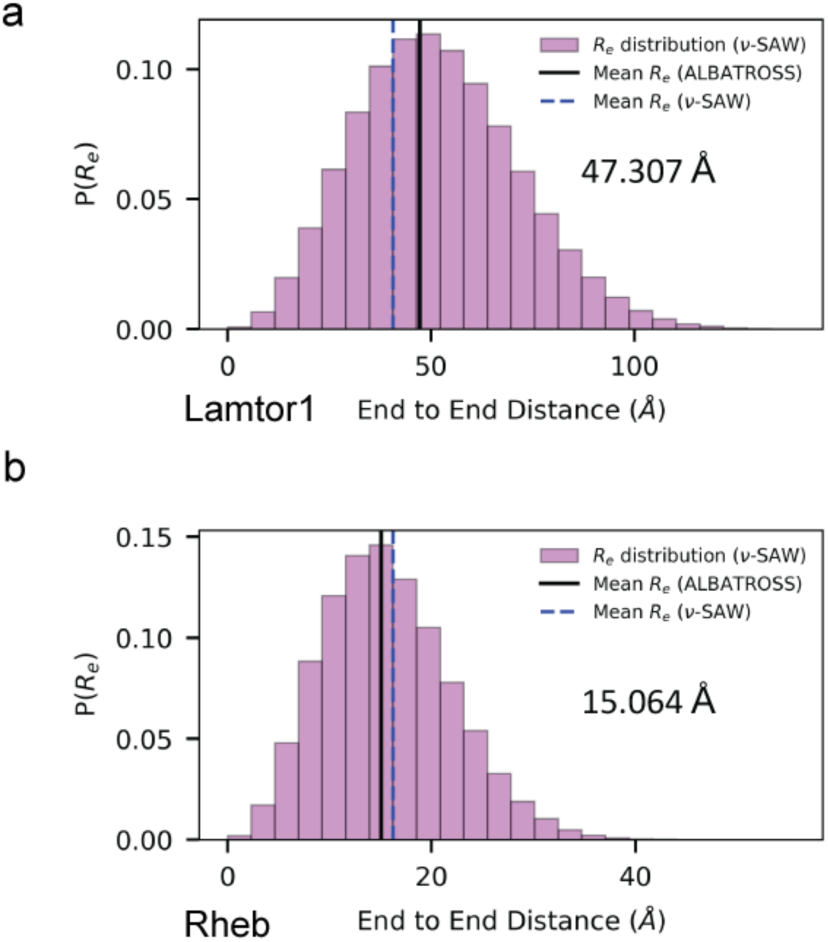
Predicted end-to-end distance for Lamtor1 and Rheb. The predictions are made by the ALBATROSS-Colab (https://colab.research.google.com/github/holehouse-lab/ALBATROSS-colab/blob/main/example_notebooks/polymer_property_predictors.ipynb). The residues from Lamtor1 (C3-A46) and Rheb (A174-S180) are used as input. The mean distance from ALBATROSS is indicated. The mean distance and distribution by *ν*-SAW model are shown with dashed blue line and pink columns, respectively.

**Extended Data Table 1.**
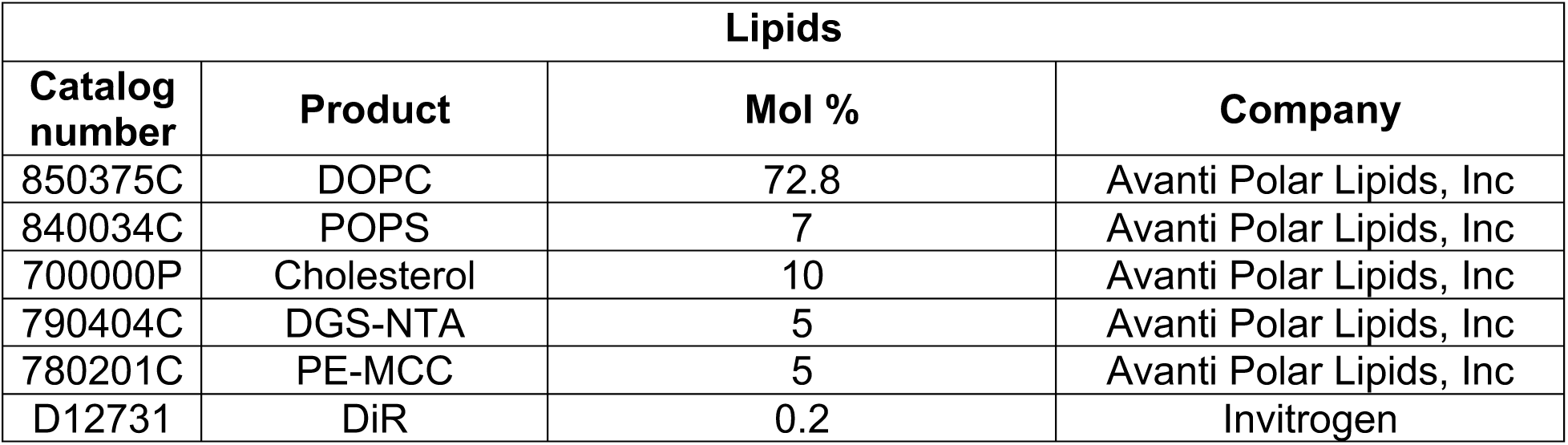
Lipid composition of liposomes used in this study.

| **Lipids** |  |  |  |
| --- | --- | --- | --- |
| **Catalog number** | **Product** | **Mol %** | **Company** |
| 850375C | DOPC | 72.8 | Avanti Polar Lipids, Inc |
| 840034C | POPS | 7 | Avanti Polar Lipids, Inc |
| 700000P | Cholesterol | 10 | Avanti Polar Lipids, Inc |
| 790404C | DGS-NTA | 5 | Avanti Polar Lipids, Inc |
| 780201C | PE-MCC | 5 | Avanti Polar Lipids, Inc |
| D12731 | DiR | 0.2 | Invitrogen |

**Extended Data Table 2.**
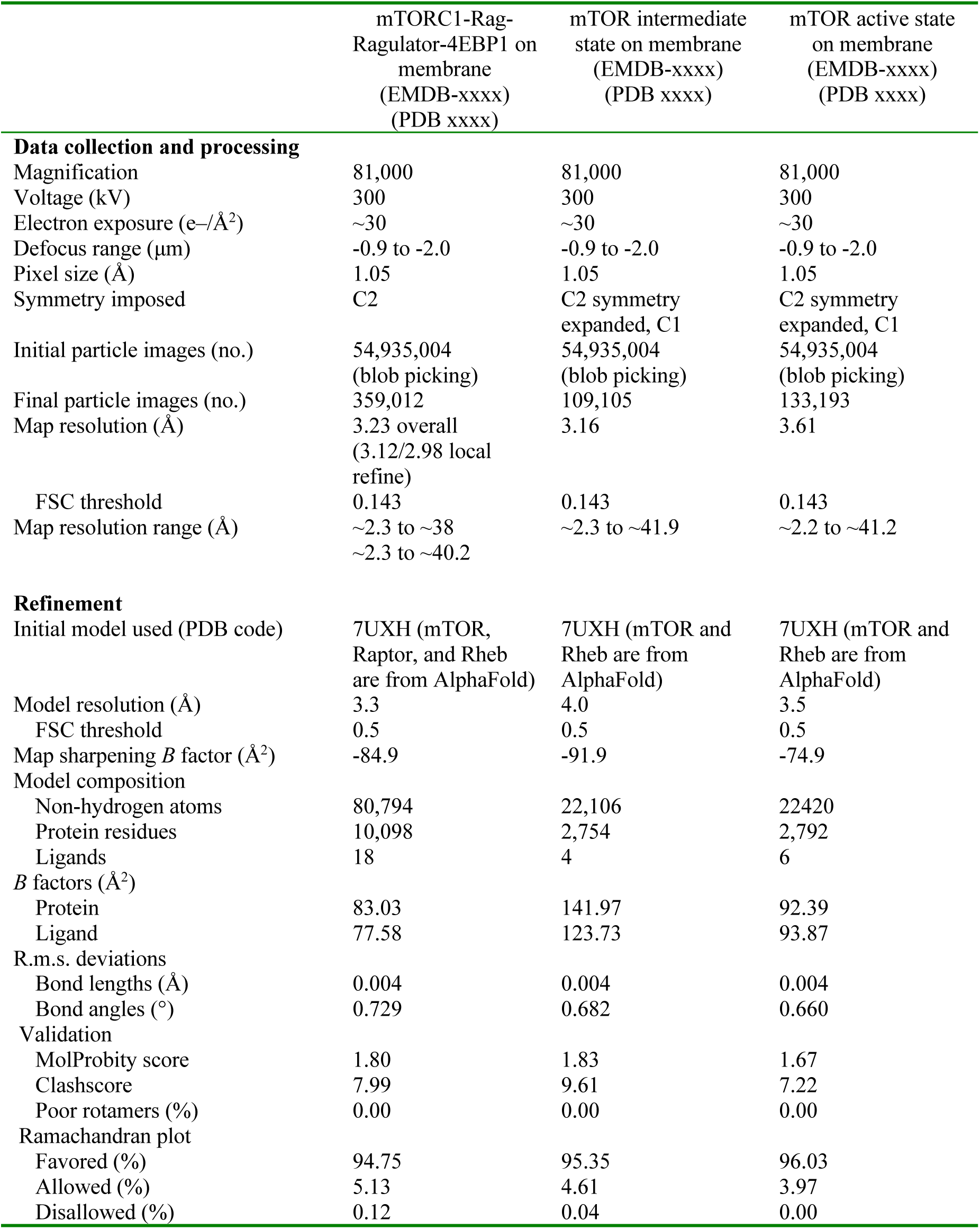
Cryo-EM data collection, refinement and validation statistics.

